# A novel mouse model of cholangiocarcinoma uncovers a role for a SOX17-Tensin 4 pathway in tumor progression

**DOI:** 10.1101/870212

**Authors:** Mickaël Di-Luoffo, Sophie Pirenne, Thoueiba Saandi, Axelle Loriot, Claude Gérard, Nicolas Dauguet, Florence Lamoline, Katarzyna Konobrocka, Vitaline De Greef, Mina Komuta, Patrick Jacquemin, Frédéric P. Lemaigre

## Abstract

**Background & Aims:** Although earlier diagnosis and treatment of intrahepatic cholangiocarcinoma (iCCA) is necessary to improve therapy, there is still limited information available about initiation and evolution of iCCA precursor lesions. Therefore, there is a need to identify mechanisms driving formation of precancerous lesions and their progression towards invasive tumor using experimental models that faithfully recapitulate human tumorigenesis.

**Methods:** We generated a new mouse model which combines cholangiocyte-specific expression of *Kras*^*G12D*^ with 3,5-diethoxycarbonyl-1,4-dihydrocollidine diet-induced inflammation to mimic iCCA development in patients with cholangitis. Histological and transcriptomic analyses of the mouse precursor lesions and iCCA were performed and compared with human analyses. The function of genes overexpressed during tumorigenesis was investigated in human cell lines.

**Results:** Mice expressing *Kras*^*G12D*^ in cholangiocytes and fed a DDC diet developed cholangitis, ductular proliferations, intraductal papillary neoplasms of bile ducts (IPNBs) and eventually iCCAs. The histology of mouse and human IPNBs were highly similar, and mouse iCCAs displayed histological characteristics of human mucin-producing large duct type iCCA. Signaling pathways activated in human iCCA were activated in mice. The identification of transition zones between IPNB and iCCA on tissue sections, combined with RNA-sequencing analyses of the lesions supported that iCCAs derive from IPNBs. We provide evidence that a gene cascade which comprises *KRAS*^*G12D*^, SRY-related HMG box transcription factor 17 (*SOX17*) and Tensin 4 (*TNS4*), and which is activated by epidermal growth factor, promotes tumor progression.

**Conclusions:** We developed a novel mouse model that faithfully recapitulates human iCCA tumorigenesis and identified a gene cascade promoting tumor progression.

## Introduction

Prognosis of intrahepatic cholangiocarcinoma (iCCA) remains dismal. Since this results in part from delayed diagnosis, there is a need to understand the mechanisms that drive development of precancerous lesions and their progression towards malignant tumor.^1^ A range of perturbed signaling pathways and genomic alterations was found to be associated with late-stage cholangiocarcinoma, but there is still limited knowledge about how such pathways and mutations evolve throughout tumorigenesis.^2-8^ Moreover, our understanding of this process is further complicated by the observation that iCCA can arise from several cell types, namely cholangiocytes, hepatocytes, progenitor cells and peribiliary glands.^9-16^

Histopathological studies identified two subtypes of iCCA in humans. Small duct subtype is typically located at the periphery of the liver and presents as tubular-type adenocarcinoma with cuboidal cells and ductular component without mucin production. The large duct subtype is predominantly found near the hilum and exhibits a tubular pattern with mucin-producing columnar cells.^17^ Histopathological studies classified biliary intraepithelial neoplasia (BilIN), intraductal tubulopapillary neoplasms and intraductal papillary neoplasms of bile ducts (IPNB) as non-invasive precursor lesions.^18-20^ Whereas BilINs develop in large bile ducts and present as flat or micropapillary dysplasia of the bile duct epithelium, IPNBs are macroscopic lesions of the large ducts characterized by exophytic papillary structures with fibrovascular core and prominent intraductal growth.^21,22^ Biliary micro-hamartomas, also called von Meyenburg complexes, are irregular ductal structures surrounded by dense stroma. These complexes are considered benign, but were reported to evolve towards iCCA in rare instances and potentially qualify as iCCA precursor lesion.^23^

Gain-of-function mutations of the small GTPase v-Ki-ras2 Kirsten rat sarcoma viral oncogene homolog (*KRAS*) are detected in iCCA at a frequency of 15-25%, and are associated with a significant decrease of survival.^24-28^ Importantly, *KRAS* mutations were detected in BilINs and IPNBs in 25-50% of cases, strongly suggesting that such mutations contribute to initiate tumorigenesis.^29-31^ Chronic inflammation is a well known risk factor of iCCA. Patients with primary sclerosing cholangitis are at high risk for iCCA and molecular classification of late-stage cholangiocarcinomas underscored the importance of inflammatory gene expression in iCCA pathogeny.^32-34^

Animal models are invaluable for investigating intra- and extrahepatic CCA development, and several models were designed based on induction of cholestasis, administration of tumor-promoting agents, injection of tumor cells, and generation of genomic alterations.^35-37^ However, limited attention was paid to tumor initiation and progression, and in several instances genetic alterations were induced at the prenatal stage or simultaneously in hepatocytes and cholangiocytes, unlike in humans. Here, we generated an original mouse model of iCCA tumorigenesis in which expression of the oncogenic mutant *Kras*^*G12D*^ is specifically induced in cholangiocytes and is combined with diet-induced inflammation that mimics chronic cholangitis. The mice developed IPNB similar to those in humans, and our data support that those lesions evolve toward malignant iCCA. We also identified a gene regulatory pathway that is activated during tumorigenesis and which promotes cell proliferation and migration.

## Material and Methods

### Animals

Osteopontin-iCreERT^2^ (Opn-Cre^ER^), loxP-stop-loxP-Kras^G12D/+^ (LSL-Kras^G12D/+^) and Rosa26R^eYFP^ mice were described^38-40^ and bred in a CD1-enriched background. For Cre-LoxP recombination, 6 week-old mice were injected intraperitoneally three times at one day interval with tamoxifen (T5648; Sigma-Aldrich, Bornem, Belgium) dissolved in corn oil at a concentration of 30 mg/mL and at a dose of 150 mg/kg of body weight. Mice were fed a standard diet or a diet containing 3,5-diethoxycarbonyl-1,4 dihydrocollidine 0.1% (DDC; 137030 Sigma-Aldrich) and monitored for signs of illness as described in the Results section and in Supplementary Figure 1. Animals received humane care according to the Directive 2010/63/EU of the European Parliament and of the Council on the protection of animals used for scientific purposes. Protocols were approved by the Animal Welfare Committee of the Université catholique de Louvain.

**Figure 1.**
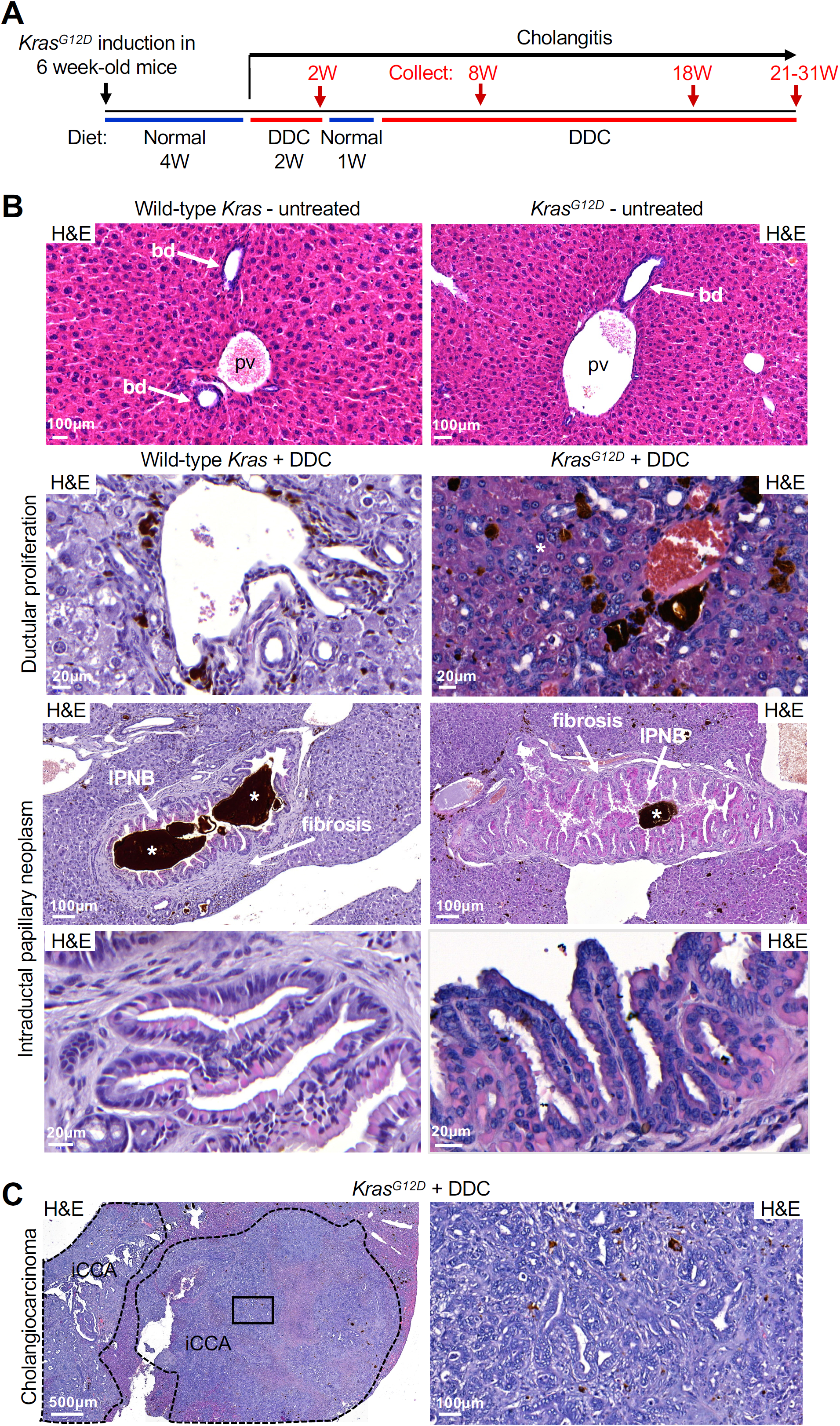
Impact of diet-induced chronic cholangitis and cholangiocyte-specific expression of *Kras*^*G12D*^ on development of biliary lesions. (*A*) Timing of *Kras*^*G12D*^ expression and of 3,5-diethoxycarbonyl-1,4-dihydrocollidine (DDC)-induced cholangitis. (*B*) Opn-Cre^ER^/LSL-Kras^G12D^ mice expressing *Kras*^*G12D*^ following tamoxifen administration (*Kras*^*G12D*^) do not show any lesion, whereas DDC-treated mice in the absence (wild-type *Kras*+DDC) or presence of KRAS^G12D^ (*Kras*^*G12D*^+DDC) develop ductular proliferations and intraductal papillary neoplasms of bile ducts (IPNB) with fibrous thickening of the bile duct wall. (*C*) Intrahepatic cholangiocarcinomas (iCCA) are detected only in the *Kras*^*G12D*^+DDC mice and starting at 21 weeks of DDC treatment. Boxed region in left panel is magnified in right panel. bd, bile duct; H&E, hematoxylin and eosin; pv: portal vein; W, weeks. *, porphyrin plug.

### Plasmids, sub-cloning and mutagenesis

*Kras*^*G12D*^, *Sox17*, and *Tns4* cDNAs were polymerase chain reaction (PCR)-amplified using the primers listed in Supplementary Table 1 and sub-cloned in pcDNA3.1+ (Thermo Fisher Scientific, Waltham, MA, USA) and pSBtet-GFP-Neomycin (pSBtet-GN; Addgene plasmid # 60501; a gift from R. Marschalek^41^). The SfiI site in mouse *Tns4* cDNA was mutated using the QuikChange II XL mutagenesis kit (Agilent, Santa Clara, CA, USA) and the primers listed in Supplementary Table 1. PCR amplification was performed using a CloneAmp™ HiFi PCR Premix (Takara, Mountain View, CA, USA) according to the manufacturer’s protocol, and amplicons were digested 30 min to 1 h at 37°C with BamHI, KpnI, XbaI or XhoI or 50°C with SfiI (NEB Ipswich, MA, USA). The ligations were performed 30 min at room temperature using a T4 DNA ligase (Thermo Fisher Scientific). pCMV(CAT)T7-SB100 plasmid coding for Sleeping Beauty Tranposase (Addgene plasmid # 34879) was a gift from Z. Izsvak.^42^

### Cell culture and transfections

Human EGI-1 cholangiocarcinoma cells (German Collection of Microorganisms and Cell Cultures, kind gift from L. Fouassier) and immortalized normal mouse cholangiocytes^43^ (NMC; kind gift from T. Shimosegawa) were grown in Dulbecco’s Modified Eagle’s Medium (DMEM; Westburg, Leusden, Netherlands), 10% fetal bovine serum (FBS; Merck, Darmstadt, Germany), 2 mM L-Glutamine (Thermo Fisher Scientific), Penicillin-Streptomycin 50 U/mL and 50 µg/mL respectively (Gibco™, Waltham, MA, USA), 2.5 µg/mL Amphotericin B (Gibco™) and 10% MEM Non-Essential Amino Acids (NEAA; Gibco™) in a humidified 5% CO_2_ incubator at 37°C. For transient transfections, cells were grown in 60 mm dishes and transfected with 3 µg plasmid, using jetPRIME® (Polyplus-Transfection, Illkirch-Graffenstaden, France) for 24 h, in at least three independent experiments. DNA was transfected 24 h after plating the cells on 60 mm dishes, and the medium was changed every day. For stable transfections, NMC or EGI-1 cells were grown in 35 mm dishes. Cells in each well were transfected with 1.9 µg plasmid (pSBtet-GN, pSBtet-GN-SOX17 or pSBtet-GN-TNS4) and 100 ng of pCMV(CAT)T7-SB100 vector, using jetPRIME® (Polyplus-Transfection, Illkirch-Graffenstaden, France). Twenty-four hours after transfection, cells were subjected to 1 mg/mL G418 for 7 days. GFP-positive cells were sorted using FACSAria III (BD, Franklin Lakes, USA) and distributed one cell per well for clonal selection. After growth for 5 days in DMEM, 10% FBS, 2 mM L-Glutamine, Penicillin-Streptomycin (50 U/mL and 50 µg/mL), 2.5 µg/mL Amphotericin B and 10% NEAA, each clone was treated with 1 mg/mL G418 for 2 weeks and the stability of the transfected vectors was monitored by detecting GFP fluorescence. Cells were treated with 0.5 µg/mL doxycycline for 5 days to induce SOX17 and TNS4.

### Laser capture microdissection (LCM), RNA extraction and sequencing

The livers were collected, embedded in OCT, frozen in liquid nitrogen and stored at −80 °C until processing. Microscope support glass, membrane slides and all tools were pre-treated to remove RNAse (RNaseZAP, Sigma-Aldrich). Ten µm sections were cut using Cryostar NX70 (Thermo Fisher Scientific) at -25°C for knife/blade temperature and -13°C for specimen temperature. Liver sections were deposited on slides adapted for LCM (SuperFrost Plus (25×75×1mm); VWR, Leuven, Belgium) and stored at -80°C inside plastic containers with silica granules to prevent tissue hydration. Within 12h, the liver sections were microdissected using Leica LMD7000 microscope (GIGA, Liège, Belgium) and RNA was directly extracted with RNAqueous®-Micro total RNA isolation AM1931 Kit (Thermo Fisher Scientific). RNA was stored at -80°C. RNA sequencing was performed by the Next Generation Sequencing Service (NGS Genewiz, South Plainfield, NJ, USA). Quality of samples was controlled by NanoDrop (NanoDrop 2000 (Thermo Fisher Scientific) and Qubit fluorometry (Invitrogen, Carlsbad, CA)) for measuring concentration and calculation of RNA integrity number (RIN). RNA was amplified using SMARTer® Universal Low Input RNA Kit for sequencing (Takara, Mountain View, CA, USA). Ribosomal RNA was depleted using RIboGone™-Mammalian rRNA removal kit (Takara, Mountain View, CA, USA), and cDNA libraries were generated using the TruSeq total stranded RNA-seq kit (Illumina, San Diego, CA, USA). cDNA libraries were sequenced using Illumina HiSeq 2×150bp configuration with estimated data output ≥350M raw paired-end reads per lane. RNA sequencing data include the generation of FASTQ-format files containing reads sequenced.

### Bioinformatics analysis workflow

Read quality control was performed using FastQC software v0.11.7 (Babraham Institute, Cambridge, UK). Low quality reads were trimmed and adapters were removed using Trimmomatic software v0.35 (RWTH Aachen University, Germany). Reads were aligned using HISAT2 software v2.1.0 (Johns Hopkins University School of Medicine, Center for Computational Biology, Baltimore, MD, USA) on GRCm38 mouse genome. Gene counts were generated using HTSeq-count (v0.5.4p3) software^44,45^ and gencode.vM15.annotation.gtf annotation file. Differential expression analysis were done using DESeq2 v1.22.2,^46^ on R version 3.5.1. Differential expression analyses were performed using adjusted p-value <0.05 and log fold-change <0.01. All differential expression data are relative data from normalized transcript per million (TPM). Pearson correlation analyses were performed using ‘rcorr’ function from the Hmisc package (RStudio).

### Cell proliferation assays

For nuclear fluorescence detection experiments, 7000 EGI-1 cells were plated at day 1 and grown for 48 h in 48-well plates. Cells were stained with Hoechst (1/5000) for 30 min and fluorescence was monitored using victor x4 2030 Multilabel Reader (PerkinElmer, Waltham, MA, USA) with excitation wavelength at 350 nm. For crystal violet staining, 4×10^4^ EGI-1 cells were plated at day 1 in 60 mm dishes and grown for minimum 3 days. At day 3, 5 and 7 cells were rinsed with PBS and fixed using 10% acetic acid/10% methanol in H_2_O for 15 min. Cells were stained with 0.5% crystal violet (#C6158, Sigma-Aldrich) in H_2_O for 15 min. The dishes were photographed and cells were counted using 16 pictures taken with help of a grid placed on each dish for each condition (Axiovert 200 fluorescent microscope 10X magnification). All pictures were transformed into white/black binary pictures and pixels were quantified using ImageJ v1.52p.

### Cell migration assay

7×10^3^ EGI-1 or 1.5×10^4^ NMC cells, wild-type or overexpressing doxycycline-induced TNS4, were plated at day 1 in culture-insert 2 wells installed in µ-Dish^35mm,high^ (#81176, Ibidi Gmgh, Gräfelfing, Gemany) and grown for 48 h. Then, the culture-inserts 2 wells were removed leaving a wound of about 200 μm between the two wells. Six adjacent pictures covering the entire wound were taken at 0h, 1h, 3h, 6h and 8h of culture. Three wound distances were measured on each picture with 50 µm interval between each measurement (ZEN Blue v2.5 software, Carl Zeiss Vision, Oberkochen, Germany).

### Statistical analyses

Single comparisons between two experimental groups were done using a paired Student’s t test. To identify significant differences between multiple groups, statistical analyses were performed using a one-way ANOVA followed by Newman-Keuls test. For adjusted p-value, a Benjamini-Hochberg correction was applied. For all analyses, p-value<0.05 was considered significant. All statistical analyses were done using the Prism software (GraphPad softwares).

### Animal housing and numbers, Western blotting, Reverse Transcription-quantitative PCR, cholangiocyte purification, tissue staining procedures

See supplementary information

## Results

### Cholangiocyte-specific expression of Kras^G12D^ combined with DDC-induced cholangitis causes stepwise development of IPNB and iCCA

To recapitulate the tumorigenic process of human iCCA, we generated a mouse model which combines cholangiocyte-specific *Kras*^*G12D*^ expression with chronic cholangitis. Opn-Cre^ER^ mice, which express tamoxifen-inducible Cre^ER^ recombinase specifically in cholangiocytes, were mated with LSL-Kras^G12D/+^ mice, which contain a mutated *Kras* allele coding for KRAS^G12D^ when a loxP-flanked cassette has been removed by CreER-mediated recombination.^38-40^ The mice also contained a tamoxifen-inducible *Rosa26R*^*eYFP*^ allele. *Kras*^*G12D*^ expression was induced by administration of tamoxifen to 6 week-old mice. Mice were then fed a normal diet for 4 weeks to enable wash out of tamoxifen. Thereafter, chronic cholangitis was induced by feeding the mice a 3,5-diethoxycarbonyl-1,4-dihydrocollidine (DDC) diet, as described by Fickert and coworkers.^47^ To this end, the mice received a DDC diet which was interrupted after 2 weeks because of weight loss. Following 1 week of weight recovery under normal diet, DDC diet was resumed, and livers were collected after 8 and 18 weeks of cholangitis, and at several time points between 21 and 31 weeks of cholangitis (Figure 1A). During this period the mice accepted the DDC diet; the experiments were interrupted when animals showed signs of severe illness and had reached humane endpoints (Supplementary Figure 1A-B).

Expression of *Kras*^*G12D*^ in the absence of DDC did not affect bile duct morphology. In contrast, administration of DDC for 8 weeks to mice which express wild-type *Kras* induced accumulation of porphyrin plugs, ductular proliferation, as well as emerging papillary lesions and periductal fibrosis of large ducts. Expression of *Kras*^*G12D*^ in mice receiving an 8-week DDC treatment was associated with large duct lesions which displayed extensive papillary proliferation, similar to IPNB (Supplementary Figure 1C).

At 18 weeks of DDC feeding, large duct lesions in mice expressing wild-type *Kras* had evolved to IPNB, and ductular proliferations were more extended (Figure 1B). Expression of *Kras*^*G12D*^ in the absence of DDC still did not affect the histology of the biliary epithelium (Figures 1B and 2A). At 18 weeks of DDC treatment, expression of *Kras*^*G12D*^ was associated with IPNBs which did not significantly differ from those monitored in mice expressing wild-type Kras (Figure 1B).

**Figure 2.**
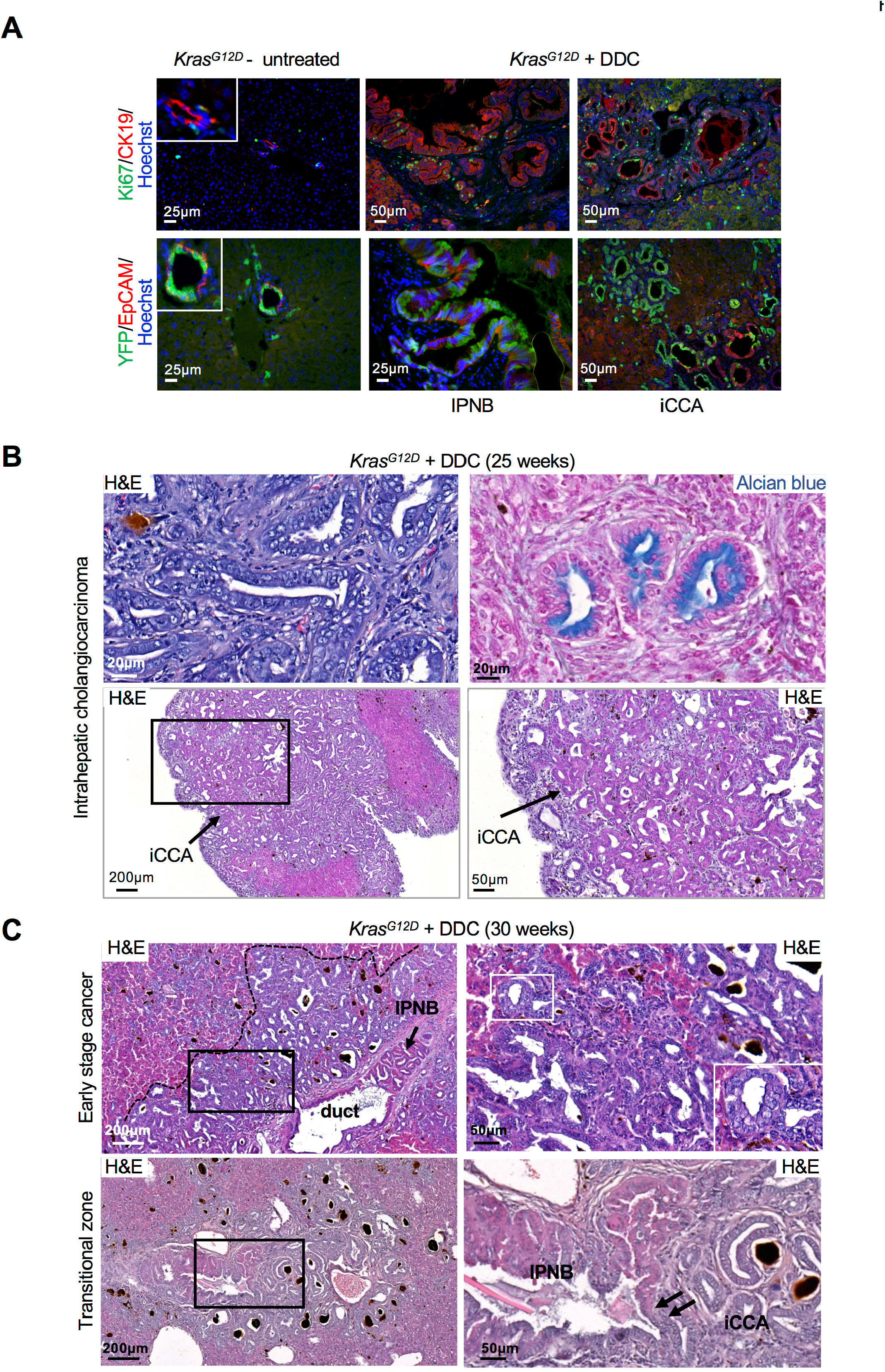
Histomorphological features of biliary lesions in mice expressing oncogenic *Kras*^*G12D*^ and fed a cholangitis-inducing diet. *(A*) In Opn-Cre^ER^/LSL-Kras^G12D^/Rosa26^eYFP^ mice fed a DDC diet for 21 weeks (*Kras*^*G12D*^+DDC), cells of IPNB and iCCA proliferate, as shown by Ki67 staining; they also express CK19 and EpCAM; eYFP expression demonstrates their cholangiocyte origin. (*B*) iCCA in *Kras*^*G12D*^+DDC mice shows irregular glandular structures displaying columnar to cuboidal morphology and fibroinflammatory stroma. iCCA cells express mucins, as evidenced by alcian blue staining. (*C*) A transition zone between IPNB and early stage of invasive iCCA (black arrows) is detected in *Kras*^*G12D*^+DDC mice. Inset in panel C (right) points to nuclear atypia. Black boxed regions in left panels are magnified in right panels.

Importantly, starting at 21 weeks of DDC diet, iCCAs were observed, but only in mice combining DDC-induced cholangitis and cholangiocyte-specific *Kras*^*G12D*^ expression (Figure 1C). iCCAs developed in one third of cases (12/36 males; 9/24 females). iCCA histology showed tubular structures with desmoplastic changes typical of human cholangiocarcinoma (Figure 1C). iCCAs were highly proliferative as illustrated by Ki67 staining, and expressed cytokeratin 19 (CK19) and epithelial cell adhesion molecule (EpCAM); they also expressed eYFP, supporting their biliary origin (Figure 2A). The epithelium was columnar to cuboidal, and was mucin-positive as shown by alcian blue staining (Figure 2B). This phenotype is similar to the human mucin-producing large duct type iCCA. Some tumors appeared less desmoplastic (Figure 2B). Interestingly, adjacent to large ducts, we often found areas of glandular proliferation with expansive growth pattern and cytonuclear atypia, which we qualified as early stage of invasive cancer (Figure 2C). Importantly, focal transitions between IPNB and early invasive cancer were detected, suggesting that IPNB evolved to iCCA (arrows in Figure 2C).

We concluded that the mouse model develops IPNB and iCCA which faithfully recapitulate the histology of their human counterparts.

### Signaling pathways known to drive iCCA development in humans are activated in the mouse model

To investigate the status of signaling pathways often perturbed in human iCCA, we first evaluated mitogen activated protein kinase (MAPK) activity and looked at phosphorylated extracellular-regulated kinase (p-ERK).^48^ p-ERK was detected in cholangiocytes following *Kras*^*G12D*^ induction or DDC treatment, and in iCCA, but not in untreated normal cholangiocytes (Figure 3A). Second, phosphoinositide-3-kinase (PI3K)^48^ was found to be activated in IPNB cells of DDC-treated mice expressing wild-type *Kras*, and in iCCA of DDC-treated mice expressing mutant *Kras*^*G12D*^, as evidenced by staining with anti-phosphorylated AKT serine/threonine kinase (p-AKT) (Figure 3A). Third, we also detected accumulation of hairy and enhancer of split-1 (HES1)^49^ in nuclei of iCCA, indicating that the Notch pathway was stimulated too (Figure 3A). Phospho-small mothers against decapentaplegic 2 (SMAD2), a marker of Transforming Growth Factor β (TGFβ) activity, is detected at high levels only in IPNB, not in other conditions. Finally, v-erb-b2 erythroblastic leukemia viral oncogene homolog 2/human epidermal growth factor receptor 2 (ERBB2/HER2)^50^ was detected in all conditions. Microdissection of IPNBs and iCCAs from tissues of mice expressing mutant *Kras*^*G12D*^ and treated for 25 weeks with DDC, followed by western blotting indicated that the amount of ERBB2 protein was higher in iCCA than in IPNB, and that ERBB2 was active, as evidenced by detection of phosphorylated ERBB2^Y1248^ (Figures 3A-B). Also, RT-qPCR analysis of total RNA from normal purified cholangiocytes and from microdissected IPNB and iCCA showed higher expression of epidermal growth factor receptor (EGFR) in the lesions (Figure 3B).

**Figure 3.**
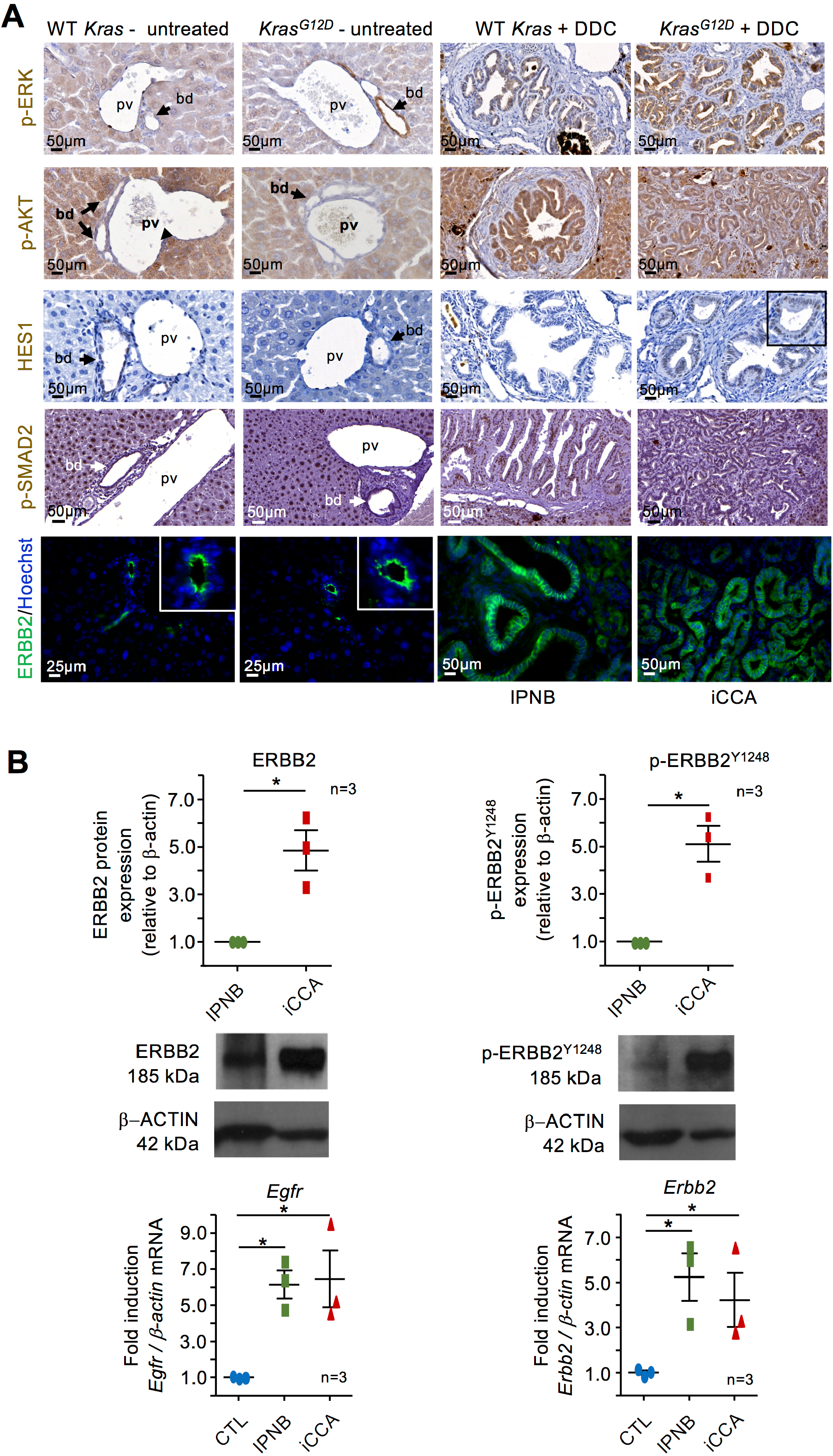
Signaling pathways activated during tumor progression in mice. (*A*) p-ERK staining is detected in cholangiocytes following *Kras*^*G12D*^ expression, in DDC-induced IPNB and *Kras*^*G12D*^+ DDC-induced iCCA. p-AKT is detected in iPNB and iCCA, while HES1 is detected only in iCCA. p-SMAD2 is detected only in IPNB. ERBB2 is detected in biliary epithelial cells in all tested conditions. (*B*) Levels of ERBB2 protein are higher in iCCA than in IPNB and ERBB2 protein is active as evidenced by detection of p-ERBB2^Y1248^. Expression of *Egfr* and *Erbb2* mRNA was higher in IPNB and iCCA than in normal purified cholangiocytes. Data in dot plots are means +/- SEM. *, p<0.05.

We concluded that MAPK, AKT, Notch, TGFβ and ERBB signaling were all activate during tumorigenesis in our mouse model, like in human iCCA.

### IPNB and iCCA display similar gene expression profiles and overexpress Sox17 and Tns4

To gain insight into the mechanisms of tumor progression we investigated the gene expression profile of ductular proliferations, IPNBs and iCCAs in DDC-treated mice expressing mutant *Kras*^*G12D*^. The three types of lesions co-existed in livers treated for 25 weeks with DDC and were isolated at that stage. Microdissected iCCAs displayed the histology of early stage of invasive cancer, as illustrated in Figure 2B. Microdissection was performed by laser capture microdissection (LCM) followed by RNA sequencing. The results were compared with those from normal cholangiocytes FACS-sorted from Opn-Cre^ER^/Rosa26R^eYFP^ mice untreated with DDC. Principal component analysis (PCA) of the RNA sequencing data identified distinct groups. Ductular proliferations and normal cholangiocytes were well separated, while iCCAs and IPNBs clustered more closely. Combined with the location of iCCAs near large ducts containing IPNB and the existence of transitional zones between IPNB and iCCA (Figure 2B), the closely related gene expression pattern suggests that iCCA originate from IPNB. Nineteen genes were found to be differentially expressed between IPNB and iCCA (fold-change >2, adjusted p-value <0.05), out of which 10 were overexpressed (Figure 4B). Except for *Aspa/ACY2*, these 10 genes were also overexpressed in human iCCA versus non-tumor tissue (The Cancer Genome Atlas cohort (TCGA); n=34; http://firebrowse.org) and/or Gene Expression Omnibus cohort GSE119336 (n=15)^51^), supporting the relevance of the mouse data with human data (Figure 4C).

**Figure 4.**
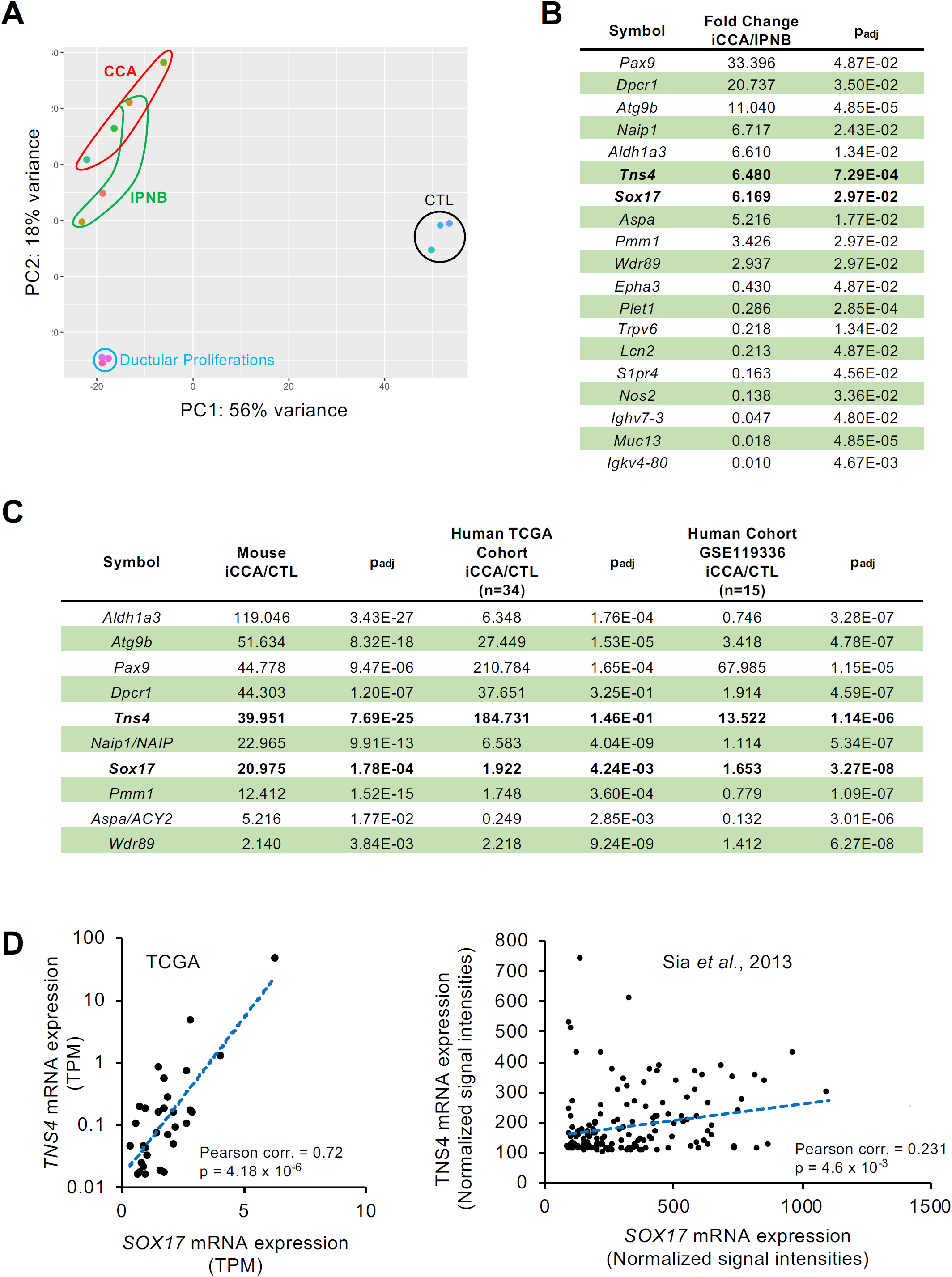
Differentially expressed genes between IPNB and iCCA. (*A*) Principal component analysis of RNA sequencing data of control cholangiocytes, ductular proliferations, IPNB and iCCA. Control cholangiocytes are purified from tamoxifen-treated Opn-Cre^ER^/Rosa26R^eYFP^ mice, the lesions were microdissected from Opn-Cre^ER^/LSL-Kras^G12D^/Rosa26R^eYFP^ mice treated with tamoxifen to induce *Kras*^*G12D*^ and fed a DDC diet for 25 weeks. (*B*) Table of the 19 genes differentially expressed between IPNB and iCCA. (*C*) Genes which are overexpressed in murine iCCA *versus* IPNB are also overexpressed in murine iCCA *versus* control cholangiocytes. Most of these genes are overexpressed in human iCCA versus control tissue. Two human iCCA cohorts are compared with mouse iCCA. (*D*) Correlation between SOX17 and TNS4 mRNA expression in human iCCA. The cohorts are described in TCGA and in Sia and coworkers.^34^

SRY-related HMG box transcription factor 17 (SOX17) and Tensin 4/C-terminal tensin-like protein (TNS4/CTEN) attracted our attention. Indeed, SOX17 cooperates with KRAS^G12D^ to promote pancreatic metaplasia and tumor progression, and the focal adhesion protein TNS4 is upregulated by EGF-ERK signaling to promote tumor progression in hepatocellular carcinoma.^52,53^ RNA sequencing and RT-qPCR showed overexpression of both genes in iCCA as compared to IPNB. Microdissection of IPNBs and iCCAs from tissues of mice expressing *Kras*^*G12D*^ and treated for 25 weeks with DDC, followed by western blotting confirmed the results for SOX17 and TNS4 at the protein level (Figure 5A). *Sox17* and *Tns4* were upregulated in iCCA as compared to control in mice as well as in humans (Figure 4C). *SOX17* and *TNS4* mRNA expression levels were correlated in human iCCA (Figure 4D, Supplementary Figure 2).

**Figure 5.**
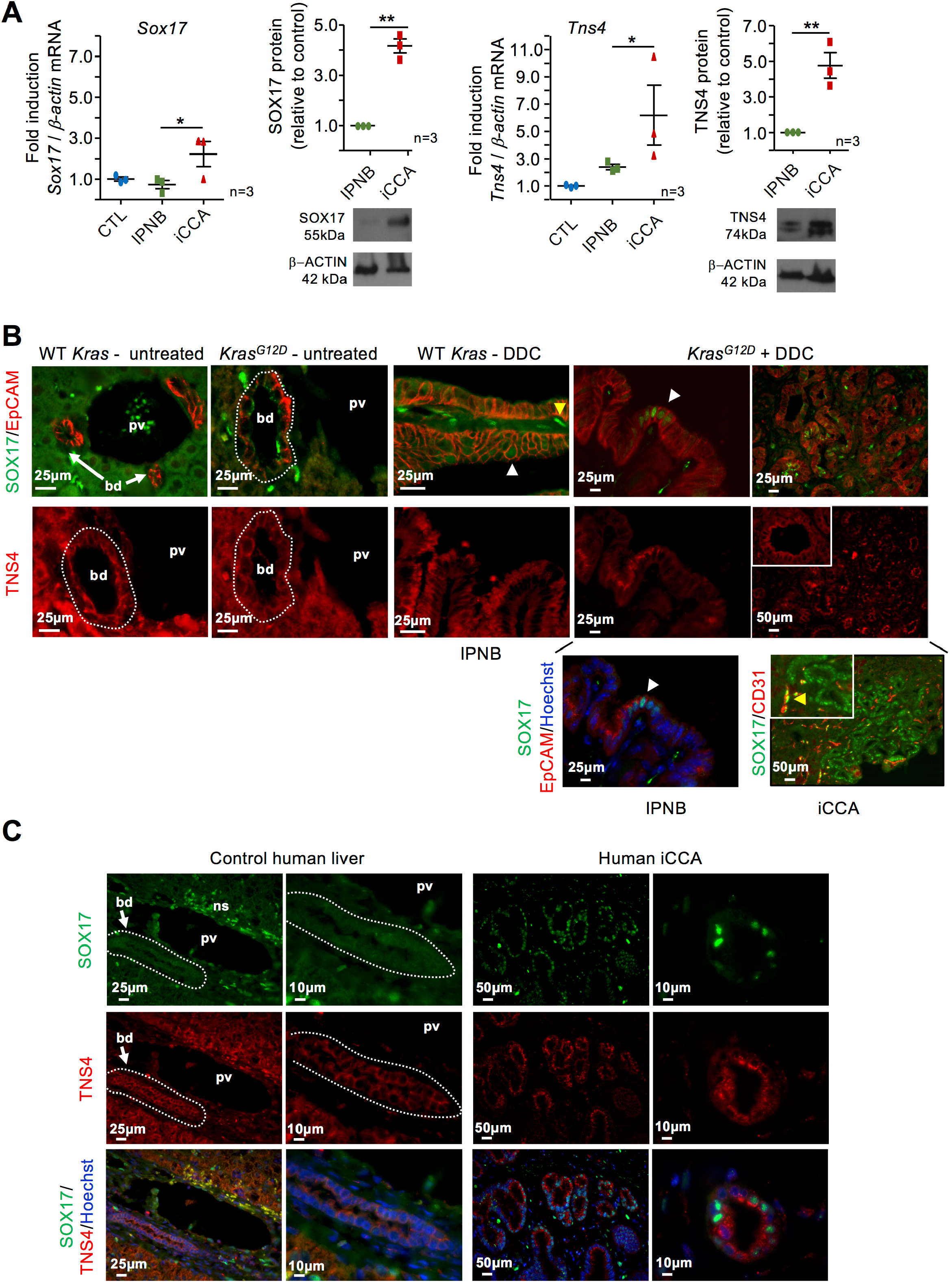
SOX17 and TNS4 expression in mouse and human iCCA. (*A*) *SOX17* and *TNS4* mRNA and protein are detected at higher levels in murine iCCA than in IPNB and control cholangiocytes. Control cholangiocytes are purified from tamoxifen-treated Opn-Cre^ER^/Rosa26R^eYFP^ mice; IPNB and iCCA are microdissected from mice expressing *Kras*^*G12D*^ and fed a DDC diet for 25 weeks. Data in dot plots are means +/- SEM. * p<0.05; ** p<0.01. (*B*) Expression of SOX17 is detected in murine DDC-induced IPNB and iCCA, not in cholangiocytes from untreated mice. TNS4 is detected in biliary cells from normal mouse ducts, IPNB and iCCA. (C) Expression of SOX17 is detected in human iCCA, not in normal cholangiocytes. Dotted lines delineate bile ducts; white arrowheads, SOX17-expressing biliary cells; yellow arrowheads, SOX17-expressing CD31-positive endothelial cells. bd, bile duct; ns, non-specific labeling; pv, portal vein. Insets represent high magnification views.

By immunostaining SOX17 was not detected in normal cholangiocytes or in *Kras*^*G12D*^-expressing cholangiocytes. In IPNBs from DDC-treated animals expressing wild-type *Kras*, a low number of cells expressed SOX17 at levels slightly higher than background. In contrast, in DDC-treated animals expressing *Kras*^*G12D*^, patches of epithelial cells in IPNB and iCCA showed high levels of nuclear SOX17 (Figure 5B). SOX17 was also detected in endothelial CD31-positive cells, as expected.^54^ Importantly, human liver displayed a similar profile with no detectable SOX17 in normal cholangiocytes, but with obvious expression in iCCA epithelial cells (Figure 5C). TNS4 was detected at the cell membrane in all tested conditions and was co-expressed with SOX17 in IPNB and iCCA (Figure 5B).

We concluded that iCCA and IPNB had similar RNA expression profiles and that SOX17 and TNS4 expression was induced in IPNB and increased in iCCA.

### SOX17 and TNS4 are induced by EGF signaling and stimulate migration and proliferation

We next explored if ERBB2 function, KRAS activation and expression of SOX17 and TNS4 were coordinated. To this end we used cholangiocarcinoma EGI-1 cells which are known to have a mutant *KRAS*^*G12D*^ gene. Treating EGI-1 cells with EGF stimulated expression of *KRAS, SOX17* and *TNS4* (Figure 6A). Similar results were found in EGF-treated NMC cells, an immortalized normal mouse cholangiocyte line that expresses high levels of cholangiocyte markers (Supplementary Figure 3A-B). Since KRAS is activated downstream of EGFR/ERBB2 and is known to stimulate SOX17 expression,^52^ we envisaged the possibility that SOX17 activates TNS4 to constitute a EGFR/ERBB2-KRAS-SOX17-TNS4 cascade (Figure 6B). We generated EGI-1 clones that stably overexpress doxycycline-inducible SOX17 or TNS4, and found that SOX17 stimulated TNS4 and to some extent *KRAS* expression, whereas TNS4 slightly induced *ERBB2* and *KRAS* (Figure 6C). Since EGI-1 cells constitutively express KRAS^G12D^, we transfected NMC cells with *Kras*^*G12D*^ to verify if it stimulates *Sox17* and *Tns4* in cholangiocytes; this was the case, further confirming the proposed gene cascade (Supplementary Figure 3C).

**Figure 6.**
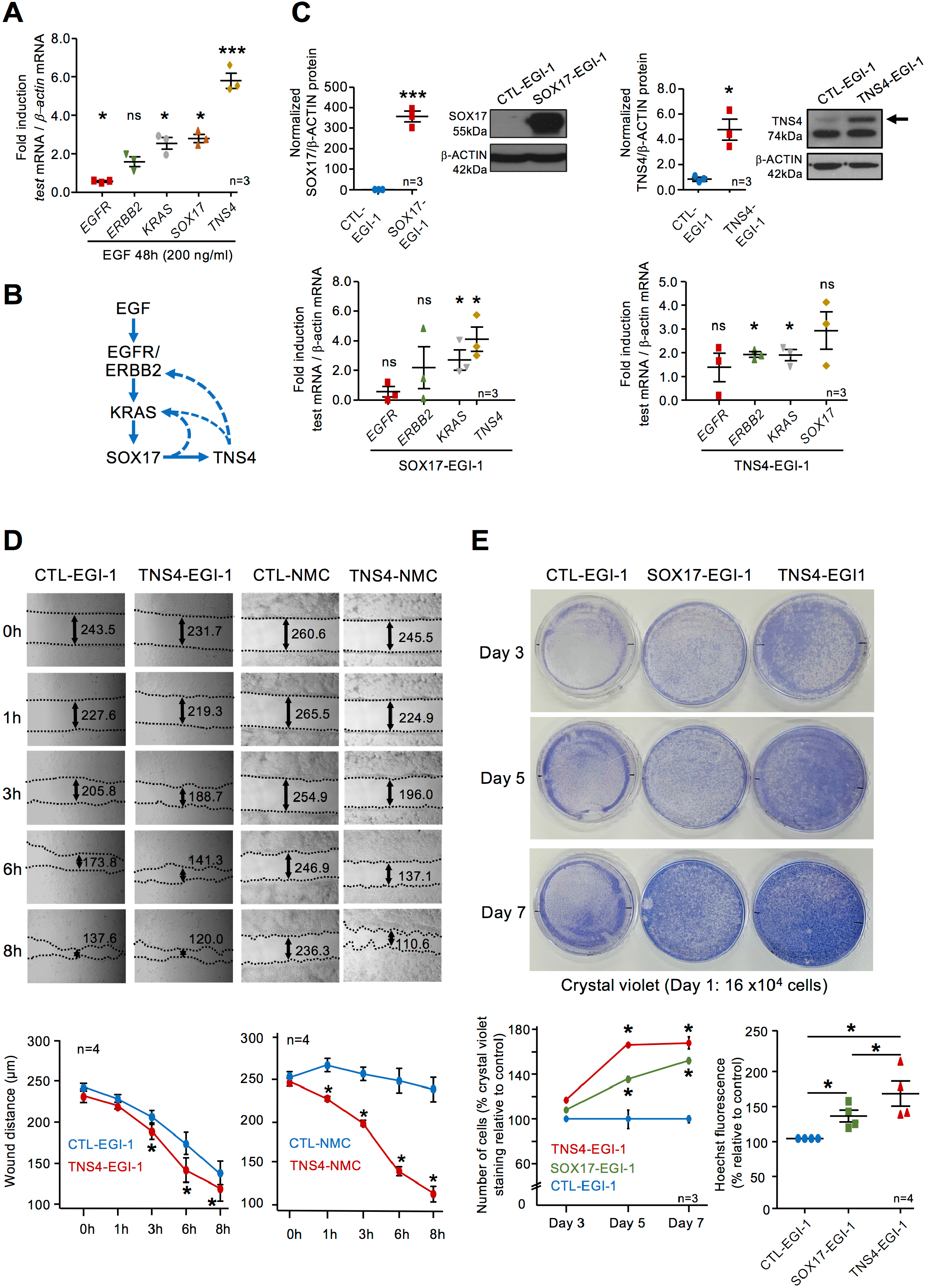
An EGF-stimulated gene cascade involving SOX17 and TNS4 is active during iCCA tumorigenesis. (*A*) Expression of *KRAS, SOX17* and *TNS4* is upregulated in EGI-1 cells incubated with EGF, as compared to untreated cells. (*B*) Structure of the EGFR/ERBB2-KRAS-SOX17-TNS4 cascade involved in iCCA tumorigenesis. (*C*) Upper panels: overexpression of SOX17 and TNS4 in EGI-1 cells stably expressing doxycycline-inducible SOX17 (SOX17-EGI-1) or TNS4 (TNS4-EGI-1), as compared to doxycycline-treated EGI-1 stably transfected with pSBtet-GN vector (CTL-EGI-1). Lower panels: SOX-17 stimulates *KRAS* and *TNS4* in SOX17-EGI-1 cells, and TNS4 stimulates *ERBB2* and *KRAS* in TNS4-EGI-1 as compared to CTL-EGI-1 cells. (*D*) Doxycycline-induced TNS4 accelerates migration of EGI-1 and NMC cells in a wound-healing assay. At each timepoint, the width of the wound is indicated as the means +/- SEM (µm) of 3 widths measured on 6 pictures covering the entire wound. (*E*) Doxycycline-induced SOX17 or TNS4 stimulates proliferation of EGI-1 cells; 16 × 10^4^ cells were plated at day 1 (upper panels). Cells illustrated in the upper panels were counted following crystal violet staining (lower left graph). In separate experiments, Hoechst fluorescence intensity of control, SOX17–EGI-1 and TNS4–EGI-1 cells was measured 4 days after plating (lower right graph). Data in graphs are represented +/- SEM. n= 3 or 4. Data in dot plot graphs represent fold inductions relative to control +/- SEM. * p<0.05 and *** p<0.001.

Since TNS4 is known to promote hepatocarcinoma cell migration,^53^ we verified if this was the case for biliary cells. Doxycycline-mediated induction of TNS4 in EGI-1 or NMC cells indeed accelerated cell migration in a wound healing assay (Figure 6D). The assay lasted for 8h, i.e. below the duration of doubling time. More important, doxycycline-mediated increase of SOX17 or TNS4 stimulated proliferation of EGI-1 cells, as shown by crystal violet staining or Hoechst fluorescence intensity assays (Figure 6E).

We concluded that a EGFR/ERBB2-KRAS-SOX17-TNS4 gene cascade is activated during iCCA tumorigenesis and contributes to cell migration and proliferation.

## Discussion

Several genetic mouse models of cholangiocarcinoma have been generated by Cre recombinase-mediated gene alterations.^35-37^ These models often induce genetic modifications at prenatal stages, or simultaneously in multiple liver cell types. For these reasons, they are not optimally suited for analysis of tumor progression starting from a well-identified adult liver cell type. Here, we selected OPN-CreER for its high efficiency and cholangiocyte specificity^38,39^ to induce expression of *Kras*^*G12D*^, a gene frequently mutated in human iCCA, including in precursor lesions. In our mouse model, *Kras*^*G12D*^ is expressed from its endogenous locus, meaning that the level of expression is within physiological limits. This was combined with chronic cholangitis in order to mimic tumor development in a context of chronic inflammation which is a major risk factor for iCCA. Thus, the cell type of origin is clearly identified, and iCCA in our mouse model develops from cholangiocytes as assessed by eYFP expression. Tumorigenesis is associated with activation of the ERK, AKT, NOTCH and ERBB pathways as in many human iCCAs. Also, the mouse iCCAs display a tubular pattern with mucin-producing columnar cells, which is typical for human large duct type iCCA; intense desmoplasia was also detected. Therefore, our animal model displays many histological and biochemical features of human iCCA. Moreover, the transcriptomic profile of IPNB and early stage of invasive iCCA, together with the histological identification of junctions between the two lesions strongly suggest that iCCA derive from IPNB. Interestingly, in humans, mutant *KRAS* is detected already in human low-grade IPNB, and tumor progression is associated with decreased TGFβ signaling,^31^ as in our model which shows strong nuclear staining of p-SMAD2 in IPNB, but no detectable p-SMAD2 in iCCA. Activation of the Wnt pathway resulting from mutations in the *Adenomatous Polyposis Coli* (*APC*) or *β-Catenin* (*CTNNB1*) gene has been reported in a subset of human IPNB,^55^ but we did not detect nuclear accumulation of β-catenin in our mouse model.

Fickert and coworkers described the inflammation occuring in DDC-treated livers.^47^ We monitored an increase in interleukin-33 (IL-33) expression in IPNB and further in iCCA (Supplementary Figure 4). IL-33 is a biliary mitogen which induces iCCA when combined with expression of constitutively active AKT and Yes-associated protein, and with obstructive cholestasis^56,57^. IL-33 is therefore likely to contribute to tumor growth in our model.

During the tumorigenic process we found that SOX17 expression in the intrahepatic cholangiocyte lineage becomes detectable at low level in IPNB of DDC-treated mice expressing wild-type *Kras*. In iCCA, SOX17 was heterogeneously distributed, with large epithelial cell patches displaying high expression of SOX17, while other tumor cells lacked detectable SOX17. The higher level of SOX17 in microdissected iCCA as compared to IPNB likely reflects the higher number of cells expressing the gene in iCCA, and possibly the fact that SOX17-positive epithelial cells in iCCA express higher levels of SOX17 than in IPNB cells, as evidenced by more intense immunostaining in iCCA than in IPNB.

SOX17 protein is not expressed in normal human or mouse intrahepatic cholangiocytes, but regulates development of the extrahepatic biliary tract and is detected in adult gallbladder epithelium.^58-61^ SOX17 is also strongly expressed in endothelial cells, which means that care is needed to interpret transcriptomic and epigenomic analyses of SOX17 expression from sampled tumor tissues composed of multiple cell types. Our data suggest that higher expression of SOX17 in iCCA results, at least in part, from EGFR/ERBB2-KRAS-mediated stimulation. SOX17 is regarded as a tumor suppressor in iCCA and downregulation of its expression by promoter hypermethylation negatively affects the patients’ prognosis.^62,63^ However, our data showing that SOX17 promotes proliferation point toward a pro-tumorigenic role. Similar to our findings, and in line with the similarity of tumorigenic processes in liver and pancreas, SOX17 was shown earlier to be induced in precursor lesions of pancreatic ductal adenocarcinoma, and to cooperate with oncogenic KRAS^G12D^ to enhance transformation of normal pancreas.^52^ Also, SOX17 is overexpressed in high grade malignant ERBB2-transformed tumor-derived cholangiocarcinoma cells as compared to less tumorigenic BDE-1 cells.^64^ The reason for the discrepancy between our data and those of Merino-Azpitarte and coworkers who showed that overexpression of SOX17 in EGI-1 cells reduces proliferation, is not clear. We note that those authors generated stable lines which overexpressed SOX17 and which displayed heterogeneous survival rates. We therefore propose that the function of SOX17 should be investigated while considering that SOX17 levels and that the heterogeneity of cellular context might determine pro-tumorigenic or tumor-suppressing properties.

Tensins are focal adhesion proteins involved in cell migration by bridging the extracellular matrix receptors and the actin skeleton. TNS4 is activated by KRAS signaling in cultured colorectal cancer cells and pancreatic adenocarcinoma cell lines in which it enhances the motility of cells.^65^ In cultured hepatocellular cancer cells TNS4 is activated by EGF-ERK signaling and also enhances cell motility.^53^ In our mouse model we detect TNS4 expression already in normal cholangiocytes, and monitor an increase in its expression during tumorigenesis. Moreover, TNS4 and SOX17 co-localized in IPNB/iCCA SOX17-positive cholangiocytes, and in transfection experiments we positioned TNS4 epistatically downstream of SOX17. TNS4 and SOX17 mRNA expression was positively correlated in human iCCA, as well as in breast carcinoma and prostate cancer but not in other tumor types (Supplementary Figure 2B)

In conclusion, we have developed a novel mouse model which recapitulates formation of iCCA precursor lesions and their evolution toward malignancy, much like in humans. The carcinogenic process is associated with activation of a EGFR/ERBB2-KRAS-SOX17-TNS4 gene cascade that contributes to cell migration and proliferation.

## Abbreviations

BilIN: biliary intraepithelial neoplasia
CK19: cytokeratin 19
DDC: 3,5-diethoxycarbonyl-1,4 dihydrocollidine
EGF: epidermal growth factor
EGFR: epidermal growth factor receptor
EpCAM: epithelial cell adhesion molecule
ERK: extracellular-regulated kinase
ERBB2: v-erb-b2 erythroblastic leukemia viral oncogene homolog 2/human epidermal growth factor receptor 2
eYFP: enhanced yellow fluorescent protein
HES1: hairy and enhancer of split-1
H&E: hematoxylin and eosin
IL-33: interleukin-33
iCCA: intrahepatic cholangiocarcinoma
IPNB: intraductal papillary neoplasm of bile ducts
KRAS: v-Ki-ras2 Kirsten rat sarcoma viral oncogene homolog
MAPK: mitogen activated protein kinase
PI3K: phosphoinositide-3-kinase
SMAD2: small mothers against decapentaplegic 2
SOX17: SRY-related HMG box transcription factor 17
TCGA: The Cancer Genome Atlas
TGFβ: transforming growth factor β
TNS4: tensin 4

## Acknowledgements

The authors thank Sandra Ormenese, Mieke Dewerchin, Javed Manesia, Satdarshan Paul Monga and Michael Oertel for advice with laser-capture microdissection; Noémie Ammar-Khodja, Jean-Nicolas Lodewijckx and Mourad El Kaddouri for help.

## References

1. Macias RIR, Kornek M, Rodrigues PM, et al. Diagnostic and prognostic biomarkers in cholangiocarcinoma. Liver Int 2019;39 Suppl 1:108–122.

2. Banales JM, Cardinale V, Carpino G, et al. Expert consensus document: Cholangiocarcinoma: current knowledge and future perspectives consensus statement from the European Network for the Study of Cholangiocarcinoma (ENS-CCA). Nat Rev Gastroenterol Hepatol 2016;13:261–80.

3. Moeini A, Sia D, Bardeesy N, et al. Molecular Pathogenesis and Targeted Therapies for Intrahepatic Cholangiocarcinoma. Clin Cancer Res 2016;22:291–300.

4. Blechacz B. Cholangiocarcinoma: Current Knowledge and New Developments. Gut Liver 2017;11:13–26.

5. Sirica AE, Gores GJ, Groopman JD, et al. Intrahepatic Cholangiocarcinoma: Continuing Challenges and Translational Advances. Hepatology 2019;69:1803–1815.

6. Labib PL, Goodchild G, Pereira SP. Molecular Pathogenesis of Cholangiocarcinoma. BMC Cancer 2019;19:185.

7. Roos E, Soer EC, Klompmaker S, et al. Crossing borders: A systematic review with quantitative analysis of genetic mutations of carcinomas of the biliary tract. Crit Rev Oncol Hematol 2019;140:8–16.

8. Fouassier L, Marzioni M, Afonso MB, et al. Signalling networks in cholangiocarcinoma: Molecular pathogenesis, targeted therapies and drug resistance. Liver Int 2019;39 Suppl 1:43–62.

9. Nakagawa H, Suzuki N, Hirata Y, et al. Biliary epithelial injury-induced regenerative response by IL-33 promotes cholangiocarcinogenesis from peribiliary glands. Proc Natl Acad Sci U S A 2017;114:E3806–E3815.

10. Guest RV, Boulter L, Kendall TJ, et al. Cell Lineage Tracing Reveals a Biliary Origin of Intrahepatic Cholangiocarcinoma. Cancer Res 2014;74:1005–1010.

11. Sekiya S, Suzuki A. Intrahepatic cholangiocarcinoma can arise from Notch-mediated conversion of hepatocytes. J Clin Invest 2012;122:3914–8.

12. Fan B, Malato Y, Calvisi DF, et al. Cholangiocarcinomas can originate from hepatocytes in mice. J Clin Invest 2012;122:2911–5.

13. Tschaharganeh DF, Xue W, Calvisi DF, et al. p53-Dependent Nestin Regulation Links Tumor Suppression to Cellular Plasticity in Liver Cancer. Cell 2016;165:1546–1547.

14. Benhamouche S, Curto M, Saotome I, et al. Nf2/Merlin controls progenitor homeostasis and tumorigenesis in the liver. Genes Dev 2010;24:1718–30.

15. Saha SK, Parachoniak CA, Ghanta KS, et al. Mutant IDH inhibits HNF-4alpha to block hepatocyte differentiation and promote biliary cancer. Nature 2014;513:110–4.

16. Komuta M, Govaere O, Vandecaveye V, et al. Histological diversity in cholangiocellular carcinoma reflects the different cholangiocyte phenotypes. Hepatology 2012;55:1876–1888.

17. Nakanuma Y, Klimstra DS, Komuta M, et al. Intrahepatic cholangiocarcinoma. WHO classification of tumours - 5th edition. Digestive system tumours: WHO, 2019:254–259.

18. Basturk O, Aishima S, Esposito I. Biliary intraepithelial neoplasia. WHO classification of tumours - 5th edition. Digestive system tumours: WHO, 2019:273–275.

19. Nakanuma Y, Basturk O, Esposito I, et al. Intraductal papillary neoplasms of the bile ducts. WHO classification of tumours - 5th edition. Digestive system tumours: WHO, 2019:279–282.

20. Kendall T, Verheij J, Gaudio E, et al. Anatomical, histomorphological and molecular classification of cholangiocarcinoma. Liver Int 2019;39 Suppl 1:7–18.

21. Aishima S, Kubo Y, Tanaka Y, et al. Histological features of precancerous and early cancerous lesions of biliary tract carcinoma. J Hepatobiliary Pancreat Sci 2014;21:448–52.

22. Ohtsuka M, Shimizu H, Kato A, et al. Intraductal papillary neoplasms of the bile duct. Int J Hepatol 2014;2014:459091.

23. Bhalla A, Mann SA, Chen S, et al. Histopathological evidence of neoplastic progression of von Meyenburg complex to intrahepatic cholangiocarcinoma. Hum Pathol 2017;67:217–224.

24. Ong CK, Subimerb C, Pairojkul C, et al. Exome sequencing of liver fluke-associated cholangiocarcinoma. Nat Genet 2012;44:690–3.

25. Churi CR, Shroff R, Wang Y, et al. Mutation profiling in cholangiocarcinoma: prognostic and therapeutic implications. PLoS One 2014;9:e115383.

26. Zou S, Li J, Zhou H, et al. Mutational landscape of intrahepatic cholangiocarcinoma. Nat Commun 2014;5:5696.

27. Nakamura H, Arai Y, Totoki Y, et al. Genomic spectra of biliary tract cancer. Nat Genet 2015;47:1003–10.

28. Javle M, Bekaii-Saab T, Jain A, et al. Biliary cancer: Utility of next-generation sequencing for clinical management. Cancer 2016;122:3838–3847.

29. Hsu M, Sasaki M, Igarashi S, et al. KRAS and GNAS mutations and p53 overexpression in biliary intraepithelial neoplasia and intrahepatic cholangiocarcinomas. Cancer 2013;119:1669–74.

30. Sasaki M, Matsubara T, Nitta T, et al. GNAS and KRAS Mutations are Common in Intraductal Papillary Neoplasms of the Bile Duct. PLoS ONE 2014;8:e81706.

31. Schlitter AM, Born D, Bettstetter M, et al. Intraductal papillary neoplasms of the bile duct: stepwise progression to carcinoma involves common molecular pathways. Mod Pathol 2014;27:73–86.

32. Khan SA, Tavolari S, Brandi G. Cholangiocarcinoma: Epidemiology and risk factors. Liver Int 2019;39 Suppl 1:19–31.

33. Andersen JB, Spee B, Blechacz BR, et al. Genomic and genetic characterization of cholangiocarcinoma identifies therapeutic targets for tyrosine kinase inhibitors. Gastroenterology 2012;142:1021–1031.

34. Sia D, Hoshida Y, Villanueva A, et al. Integrative molecular analysis of intrahepatic cholangiocarcinoma reveals 2 classes that have different outcomes. Gastroenterology 2013;144:829–40.

35. Loeuillard E, Fischbach SR, Gores GJ, et al. Animal models of cholangiocarcinoma. Biochim Biophys Acta 2018;1865:982–992.

36. Cadamuro M, Brivio S, Stecca T, et al. Animal models of cholangiocarcinoma: What they teach us about the human disease. Clin Res Hepatol Gastroenterol 2018;42:403–415.

37. Pirenne S, Lemaigre F. Genetically-engineered animal models of biliary tract cancers. Current Opinion in Gastroenterology 2020;220, in press

38. Espanol-Suner R, Carpentier R, Van Hul N, et al. Liver progenitor cells yield functional hepatocytes in response to chronic liver injury in mice. Gastroenterology 2012;143:1564–1575.

39. Lesaffer B, Verboven E, Van Huffel L, et al. Comparison of the Opn-CreER and Ck19-CreER Drivers in Bile Ducts of Normal and Injured Mouse Livers. Cells 2019;8:380.

40. Hingorani SR, Petricoin EF, Maitra A, et al. Preinvasive and invasive ductal pancreatic cancer and its early detection in the mouse. Cancer Cell 2003;4:437–50.

41. Kowarz E, Loscher D, Marschalek R. Optimized Sleeping Beauty transposons rapidly generate stable transgenic cell lines. Biotechnol J 2015;10:647–53.

42. Mates L, Chuah MK, Belay E, et al. Molecular evolution of a novel hyperactive Sleeping Beauty transposase enables robust stable gene transfer in vertebrates. Nat Genet 2009;41:753–61.

43. Ueno Y, Alpini G, Yahagi K, et al. Evaluation of differential gene expression by microarray analysis in small and large cholangiocytes isolated from normal mice. Liver Int 2003;23:449–59.

44. Anders S, Pyl PT, Huber W. HTSeq--a Python framework to work with high-throughput sequencing data. Bioinformatics 2015;31:166–9.

45. von Ahlfen S, Missel A, Bendrat K, et al. Determinants of RNA quality from FFPE samples. PLoS One 2007;2:e1261.

46. Anders S, Huber W. Differential expression analysis for sequence count data. Genome Biol 2010;11:R106.

47. Fickert P, Stoger U, Fuchsbichler A, et al. A new xenobiotic-induced mouse model of sclerosing cholangitis and biliary fibrosis. Am J Pathol 2007;171:525–36.

48. Schmitz KJ, Lang H, Wohlschlaeger J, et al. AKT and ERK1/2 signaling in intrahepatic cholangiocarcinoma. World J Gastroenterol 2007;13:6470–7.

49. El Khatib M, Bozko P, Palagani V, et al. Activation of Notch signaling is required for cholangiocarcinoma progression and is enhanced by inactivation of p53 in vivo. PLoS One 2013;8:e77433.

50. Sirica AE. Role of ErbB family receptor tyrosine kinases in intrahepatic cholangiocarcinoma. World J Gastroenterol 2008;14:7033–58.

51. Zhou G, Cao P, Li Y. RNA over-editing leads to aggressiveness of intrahepatic cholangiocarcinoma. Gene Expression Omnibus, accession number GSE119336 2019.

52. Delgiorno KE, Hall JC, Takeuchi KK, et al. Identification and manipulation of biliary metaplasia in pancreatic tumors. Gastroenterology 2014;146:233–244

53. Chan LK, Chiu YT, Sze KM, et al. Tensin4 is up-regulated by EGF-induced ERK1/2 activity and promotes cell proliferation and migration in hepatocellular carcinoma. Oncotarget 2015;6:20964–76.

54. Corada M, Orsenigo F, Morini MF, et al. Sox17 is indispensable for acquisition and maintenance of arterial identity. Nat Commun 2013;4:2609.

55. Fujikura K, Akita M, Ajiki T, et al. Recurrent Mutations in APC and CTNNB1 and Activated Wnt/beta-catenin Signaling in Intraductal Papillary Neoplasms of the Bile Duct: A Whole Exome Sequencing Study. Am J Surg Pathol 2018;42:1674–1685.

56. Li J, Razumilava N, Gores GJ, et al. Biliary repair and carcinogenesis are mediated by IL-33-dependent cholangiocyte proliferation. J Clin Invest 2014;124:3241–51.

57. Yamada D, Rizvi S, Razumilava N, et al. IL-33 facilitates oncogene-induced cholangiocarcinoma in mice by an interleukin-6-sensitive mechanism. Hepatology 2015;61:1627–42.

58. Uemura M, Ozawa A, Nagata T, et al. Sox17 haploinsufficiency results in perinatal biliary atresia and hepatitis in C57BL/6 background mice. Development 2013;140:639–48.

59. Spence JR, Lange AW, Lin SC, et al. Sox17 regulates organ lineage segregation of ventral foregut progenitor cells. Dev Cell 2009;17:62–74.

60. Higashiyama H, Ozawa A, Sumitomo H, et al. Embryonic cholecystitis and defective gallbladder contraction in the Sox17-haploinsufficient mouse model of biliary atresia. Development 2017;144:1906–1917.

61. Uhlen M, Oksvold P, Fagerberg L, et al. Towards a knowledge-based Human Protein Atlas. Nat Biotechnol 2010;28:1248–50.

62. Merino-Azpitarte M, Lozano E, Perugorria MJ, et al. SOX17 regulates cholangiocyte differentiation and acts as a tumor suppressor in cholangiocarcinoma. J Hepatol 2017;67:72–83.

63. Goeppert B, Konermann C, Schmidt CR, et al. Global alterations of DNA methylation in cholangiocarcinoma target the Wnt signaling pathway. Hepatology 2014;59:544–54.

64. Dumur CI, Campbell DJ, DeWitt JL, et al. Differential gene expression profiling of cultured neu-transformed versus spontaneously-transformed rat cholangiocytes and of corresponding cholangiocarcinomas. Exp Mol Pathol 2010;89:227–35.

65. Al-Ghamdi S, Albasri A, Cachat J, et al. Cten is targeted by Kras signalling to regulate cell motility in the colon and pancreas. PLoS One 2011;6:e20919.

