## Supplementary information for "A novel mouse model of cholangiocarcinoma uncovers a role for a SOX17-Tensin 4 pathway in tumor progression"

#### Supplementary Materials and Methods

##### *Animals*

All mice were maintained at 23°C +/- 2°C with a 12 h day/night cycle, in individually ventilated cages (IVC). Sterilized food and bedding, and acidified water were provided. Immunodeficient sentinels were exposed to dirty bedding from other cages belonging the same IVC rack. Sanitary screenings of the sentinels were performed quarterly according to FELASA guidelines; animals were free of pathogens.

Animals were monitored daily and received humane care according to the Directive 2010/63/EU of the European Parliament and of the Council on the protection of animals used for scientific purposes; the Belgian law of May 29, 2013 on protection of animals used for experimentation, updated on September 7, 2017 by the Brussels Region.

Protocols were approved by the Animal Welfare Committee of the Université catholique de Louvain with license number 2018/UCL/MD/014.
Animal numbers: Wild-type mice or tamoxifen-treated OPN-CreER mice (devoid of mutated *loxP-stop-loxP-Kras*$^{G12D}$ allele) were indistinguishable at the histological level and were used as controls. The number of mice used in the study was as follows: wild-type *KRas* + DDC: 20; *Kras*$^{G12D}$: 30; *Kras*$^{G12D}$+DDC: 65.

##### *Protein extraction and western blotting*

Cells were rinsed twice with phosphate-buffered saline (PBS) and total proteins were extracted using cell lysis buffer (50 mM Tris-Cl pH 7.4, 150 mM NaCl, 1 mM EDTA, 0.5% Igepal, 1 mM dithiothreitol and protease/phosphatase inhibitors (Roche, Bâle, Switzerland). Cells were sonicated for 15 sec and centrifuged. Protein concentration was measured using a Bradford assay. Proteins (40 µg) were fractionated using 8-10% polyacrylamide gels, transferred to PVDF membranes (Merck, Darmstadt, Germany), and detected by enhanced chemiluminescence (SuperSignal™ West Pico Plus, Thermo Fisher Scientific) using X-ray films (Thermo Fisher Scientific) and Fusion Solo S equipment (Vilber Lourmat, Collegien, France). Proteins were quantified using Bio1D advanced software (Vilber Lourmat). Primary and secondary antibodies are described in Supplementary Tables 2 and 3.

##### *RNA extraction and Reverse Transcription-quantitative PCR*

RNA was extracted from mouse tissues using RNAqueous™-Micro Total RNA Isolation Kit (Thermo Fisher Scientific) according to the manufacturer's protocol. Total RNA from cultured cells were extracted using TriPure Isolation reagent (Roche) according to the manufacturer's protocol. cDNA synthesis was performed with MMLV reverse transcriptase (Thermo Fisher Scientific) according to manufacturer's protocol. Gene expression was quantified by qPCR using Kappa Syberfast qPCR kit (Sopachem, Nazareth, Belgium) using the following conditions: 3 min at 95°C, 40 cycles of denaturation (3 sec at 95°C), annealing and extension (30 sec at 60°C). β-actin was used as internal control. Oligonucleotides used in qPCR are described in Supplementary Table 1. Quantification of gene expression was performed using the 2(-Delta Delta C(T)) method[44] and was expressed as a ratio of target to β-actin levels.

Each amplification was performed in duplicate using minimum three distinct RNA samples.

##### *Histological examination*

All tissue sections were analysed by at least two independent investigators.

*Tissue staining*

Hematoxylin/Eosin (H&E) staining was performed on 6 µm sections of formalin-fixed paraffin-embedded tissues. Briefly, tissue sections were deparaffinized 3x 3 min in xylene, 3 min in 99%, 95%, 70% and 30% ethanol and deionized $H_2O$. The sections were stained 7 sec in 100% hematoxylin, rinsed with $H_2O$, stained 7 sec in eosine, and rinsed with deionized $H_2O$. Dehydration of sections was performed in deionized $H_2O$, followed by 30%, 70%, 95%, 99% ethanol for 30 sec, and 30 sec in xylene. Coverslips were placed on slides using Depex mounting medium (VWR, Leuven, Belgium). Pictures were taken with panoramic P250 Digital Slide Scanner (Histogenex, Antwerpen, Belgium) using 3DHISTECH Case Viewer software.

*Immunohistochemistry and Immunofluorescence*

Immunohistochemistry and immunofluorescence were performed on 6 µm sections of formalin-fixed paraffin-embedded tissues. Briefly, tissue sections were deparaffinized as described above, followed by antigen unmasking by micro-wave heating for 10 min or incubation in PT Module (Lab Vision, Westinghouse Drive Fremont, CA ,USA) for 20 min at 98°C in 10 mM sodium citrate pH 6.0 or EDTA pH 8.0. Sections were permeabilized for 5 min in 0.3 % Triton X-100 PBS solution before blocking for 45 min in 0.3 % milk,10 % bovine serum albumin, 0.3 % Triton X-100 in PBS. Primary and secondary antibodies were diluted in blocking solution and incubated respectively at 4°C overnight and 37°C for 1 h. Pictures for immunohistochemistry and immunofluorescence were performed with panoramic P250 Digital Slide Scanner (Histogenex) using 3DHISTECH Case Viewer software, and Axiovert 200 fluorescent microscope and AxioVision system. Primary and secondary antibodies are described in Supplementary Tables 2 and 3.

##### *Cholangiocyte purification*

The tissular dissociation protocol was adapted from Assi et al.[48] Livers were injected with 5 mL of 600 µg/mL collagenase P solution in EGTA-buffer (0.1 M EGTA, 125 mM NaCl, 0.8 mM MgSO4, 5.4 mM KCl, 1 mM NaH2PO4, 4.2 mM NaHCO3, 10 mM HEPES and 5.6 mM glucose), collected and rinsed using PBS, and incubated in 20 mL of the same collagenase P in EDTA-buffer solution during 10 min at 37°C with agitation. Then, livers were cut in small fractions and incubated in 600 µg/mL collagenase P solution in CaCl2-buffer (1.8 mM CaCl2, 123 mM NaCl, 0.8 mM MgSO4, 5.4 mM KCl, 1 mM NaH2PO4, 4.2 mM NaHCO3, 10 mM HEPES, and 5.6 mM glucose) during 10 min at 37°C with agitation. Thereafter, individual cells were collected using a 40 µm filter and centrifuged at 1000 rpm during 5 min. Incubation with collagenase P solution in CaCl2-buffer, 40 µm filter cell filtration and centrifugation was repeated 3 times. Cells were pooled and cell pellets were rinsed 3 times with PBS and resuspended in 15 mL PBS-EDTA (1 mM). To eliminate red blood cells, cells were incubated in 3 mL of Red Blood Cell lysis buffer (Thermo Fisher Scientific) during 3 min at room temperature. The lysis reaction was quenched with 10 mL of PBS-EDTA (1mM) - FBS 1% during 1 min and centrifuged at 1000 rpm during 5 min. The cell pellets were resuspended in 2 mL of PBS-EDTA (1 mM) - FBS 1% with RNaseOUT™ Recombinant Ribonuclease Inhibitor (2000 U/mL) (Thermo Fisher Scientific). Then, GFP-positive cells were sorted using (FACSAria III), harvested in individual microtubes and centrifuged at 1000 rpm during 5 min. The supernatant was removed and RNA from cell pellets was extracted using RNAqueous™-Micro Total RNA Isolation Kit (Thermo Fisher Scientific) according to the manufacturer’s protocol.

### Supplementary Tables

#### *Supplementary Table 1. List of primers*

| APPLICATION | GENE | SEQUENCE |
|---|---|---|
| cDNAs, PCR | *Sox17* | Forward: 5'-AAAggccTCTGAggccGCGCCATGAGCAGCCCGGATG-3' |
| | | Reverse: 5'-AAAggccTGACAggccCGGATCAGGGACCTGTCACACGTCAGGATAG-3' |
| | *Tns4* | Forward: 5'- AAAggccTCTGAggccCCCATCATGTCCAGCTCTC-3' |
| | | Reverse: 5'-AAAggccTGACAggccGGCACCCTTCTACATCCTTTC-3‘ |
| Mutagenesis | *Tns4* | Forward: 5'-CTGGGGAGCAGGCaTCAGGGGCATCGCaGGATCTGG-3' |
| | | Reverse: 5'-CCAGATCCTGCGAtGCCCCTGAtGCCTGCTCCCCAG-3' |
| RT-qPCR | *β-actin* | Forward: 5'-TCCTGAGCGCAAGTACTCTGT-3' |
| Mus musculus | | Reverse: 5'-CTGATCCACATCTGCTGGAAG-3' |
| | *Ck19* | Forward: 5'-ACCCTCCCGAGATTACAACC-3' |
| | | Reverse: 5'-GGCGAGCATTGTCAATCTGT-3' |
| | *Egfr* | Forward: 5'-GCCATCTGGGCCAAAGATACC-3' |
| | | Reverse: 5'-GTCTTCGCATGAATAGGCCAA-3' |
| | *Epcam* | Forward: 5'-CTGGGAGGAGGATAAAGC-3' |
| | | Reverse: 5'-AGAAGAATGGAACAGGGAC-3' |
| | *Erbb2* | Forward: 5'-TAACTGGACCCCAGCCTATG-3' |
| | | Reverse: 5'-AACGGAGAATGACCCTGTTG-3' |
| | *Hnf1*β | Forward: 5'-GAAAGCAACGGGAGATCCTC-3' |
| | | Reverse: 5'-GACTGCCCAGGCCCCTGGTTCTGT-3' |
| | *Kras* | Forward: 5’-ACAGGCTCAGGAGTTAGCAAGGA-3’ |
| | | Reverse: 5’-AAGGCATCGTCAACACCCTGTC-3’ |
| | *Sox9* | Forward: 5’-CAAGACTCTGGGCAAGCTCTG-3’ |
| | | Reverse: 5’-TCCGCTTGTCCGTTCTTCAC-3’ |
| | *Sox17* | Forward: 5’-GAGCAGCACCTCCAGACAT-3’ |
| | | Reverse: 5’-CTTCATGCGCTTCACCTGCTT-3’ |
| | *Tns4* | Forward: 5’-GTTGCTTCCCTCTGGGACTG-3’ |
| | | Reverse: 5’-GAGAGGAAGAGGCTGCCAC-3’ |
| RT-qPCR | *β-ACTIN* | Forward: 5'-TCCTGAGCGCAAGTACTCTGT-3' |
| Homo sapiens | | Reverse: 5'-CTGATCCACATCTGCTGGAAG-3' |
| | *EGFR* | Forward: 5’-GCCCTCAACACAGTGGAGC-3’ |
| | | Reverse: 5’-CCAGCAGCTCCCATTGGG-3’ |
| | *ERBB2* | Forward: 5'-CTGCGCAGGCAGTGATGA-3' |
| | | Reverse: 5'-GTAGCCCTGCACCTCCTG-3' |
| | *KRAS* | Forward: 5'-CTGCTGCAGACAGTGAGTC-3' |
| | | Reverse: 5'-CTTGCTAAGTCCTGAGCCTG-3' |
| | *SOX17* | Forward: 5'-GCAGAATCCAGACCTGCAC-3' |
| | | Reverse: 5'-TTCAGCCGCTTCACCTGC-3' |
| | *TNS4* | Forward: 5'-CCCCCAGCATCTCAATCCC-3' |
| | | Reverse: 5'-CTGGAGTGCAGGAGGGAC-3' |

#### *Supplementary Table 2. Primary antibodies*

| PRIMARY ANTIBODY | SPECIES | SOURCE | REFERENCE NUMBER | DILUTION IHC/IF | DILUTION WB |
|---|---|---|---|---|---|
| CD31 | Rat (IgG2a) | BD Biosciences | #550274 | 1/50 | |
| CK19 | Rat | DSHB | #TROMAIII | 1/100 | |
| EGFR | Rabbit | Cell Signaling | #11/2017 | 1/50 | |
| EPCAM | Rabbit | Abcam | #ab32392 | 1/250 | |
| ERBB2 | Rabbit | Dako | #A0485 | 1/300 | 1/1000 |
| GFP | Goat | Abcam | #ab6673 | 1/250 | |
| HES1 | Rabbit | Cell Signaling | #11988S | 1/100 | |
| Ki67 | Mouse | BD Biosciences | #556003 | 1/250 | |
| p-AKT(Thr308) | Rabbit | Cell Signaling | #9275S | 1/200 | |
| p-ERBB2(Tyr1248) | Rabbit | R&D Systems | #AF1768 | | 1/1000 |
| p-ERK | Rabbit | Cell Signaling | #4370S | 1/200 | |
| p-Smad2 (S465/467) | Rabbit | Merck | #AB3849 | 1/200 | |
| SOX17 | Goat | R&D Systems | #AF1924 | 1/100 | 1/500 |
| TNS4 | Mouse | R&D Systems | #MAB6925 | 1/100 | 1/500 |
| β-ACTIN | Mouse | Sigma Aldricht | #A5441 | | 1/10000 |

#### *Supplementary Table 3. Secondary antibodies*

| SECONDARY ANTIBODY | SPECIES | SOURCE | REFERENCE NUMBER | DILUTION IHC | DILUTION WB |
|---|---|---|---|---|---|
| Alexa Fluor 488 anti-goat | Donkey | ThermoFisher Scientific | #A-11055 | 1/1000 | |
| Alexa Fluor 488 anti-mouse | Donkey | ThermoFisher Scientific | #A-21202 | 1/1000 | |
| Alexa Fluor 488 anti-rabbit | Donkey | ThermoFisher Scientific | #A-21206 | 1/1000 | |
| Alexa Fluor 594 anti-mouse | Donkey | ThermoFisher Scientific | #A-21203 | 1/1000 | |
| Alexa Fluor 594 anti-rabbit | Donkey | ThermoFisher Scientific | #A-21207 | 1/1000 | |
| Alexa Fluor 594 anti-rat | Donkey | ThermoFisher Scientific | #A-21209 | 1/1000 | |
| Anti-goat HRP-linked | Donkey | ThermoFisher Scientific | PA1-28664 | 1/500 | 1/5000 |
| Anti-mouse IgG HRP-linked | Mouse | Cell Signaling | #7076S | | 1/5000 |
| Anti-rabbit biotinylated | Sheep | Bethyl | #A120-100B | 1/1000 | |
| Anti-rabbit HRP-linked | Goat | Enzo | #ADI-SAB-300-J | | 1/5000 |
| Anti-rabbit HRP-linked | Goat | ThermoFisher Scientific | #31460 | 1/1000 | |

#### Supplementary Figure Legends

**Supplementary Figure 1**. (*A*) Time-course of mouse body weight progression during DDC treatment. Arrows point to the timing of liver collection after initiation of DDC treatment. Data are means +/- SEM. ***(B)*** Hepatomegaly in DDC-treated mice. The liver of mice expressing *Kras*$^{G12D}$ and fed a DDC diet for 25 weeks is illustrated. (*C*) Untreated mice expressing *Kras*$^{G12D}$ in cholangiocytes have normal ducts. After 8 weeks of DDC treatment, mice expressing wild type *Kras* showed emerging papillary lesions of large bile ducts, whereas DDC-treated mice expressing *Kras*$^{G12D}$ in cholangiocytes displayed IPNBs.

**Supplementary Figure 2. (***A*) Correlation between *SOX17* and *TNS4* mRNA expression in the GSE119336 cohort. The correlation is not statistically significant in iCCA and the number of samples is low, but clearly differs from the controls. Control is non-tumor adjacent tissue. (*B*) Correlation between *SOX17* and *TNS4* mRNA expression in TCGA cohorts. For each cohort from TCGA, we converted the "scaled_estimate" in the "illuminahiseq_rnaseqv2_unc_edu_Level_3_RSEM_genes" file into TPM by multiplying by $10^6$.

**Supplementary Figure 3.** (*A*) Expression of *Kras*, *Sox17* and *Tns4* is upregulated in NMC cells incubated with EGF, as compared to untreated cells. (*B*) NMC cells express cholangiocyte markers. The level of marker expression in cholangiocytes is compared with that of total adult mouse liver. (*C*) Transient transfection of *Kras*$^{G12D}$ stimulates *Sox17* and *Tns4* expression in NMC cells.

**Supplementary Figure 4.** Expression of IL-33 mRNA in control cholangiocytes purified from tamoxifen-treated Opn-Cre$^{ER}$/ROSA26R$^{eYFP}$ mice and from ductular proliferations (Duct Prol), IPNB and iCCA from Opn-Cre$^{ER}$/LSL-Kras$^{G12D}$ mice treated with tamoxifen to induce *Kras*$^{G12D}$ and fed a DDC diet for 25 weeks. Data are means of fold inductions +/- SEM.

Supplementary Figure 1

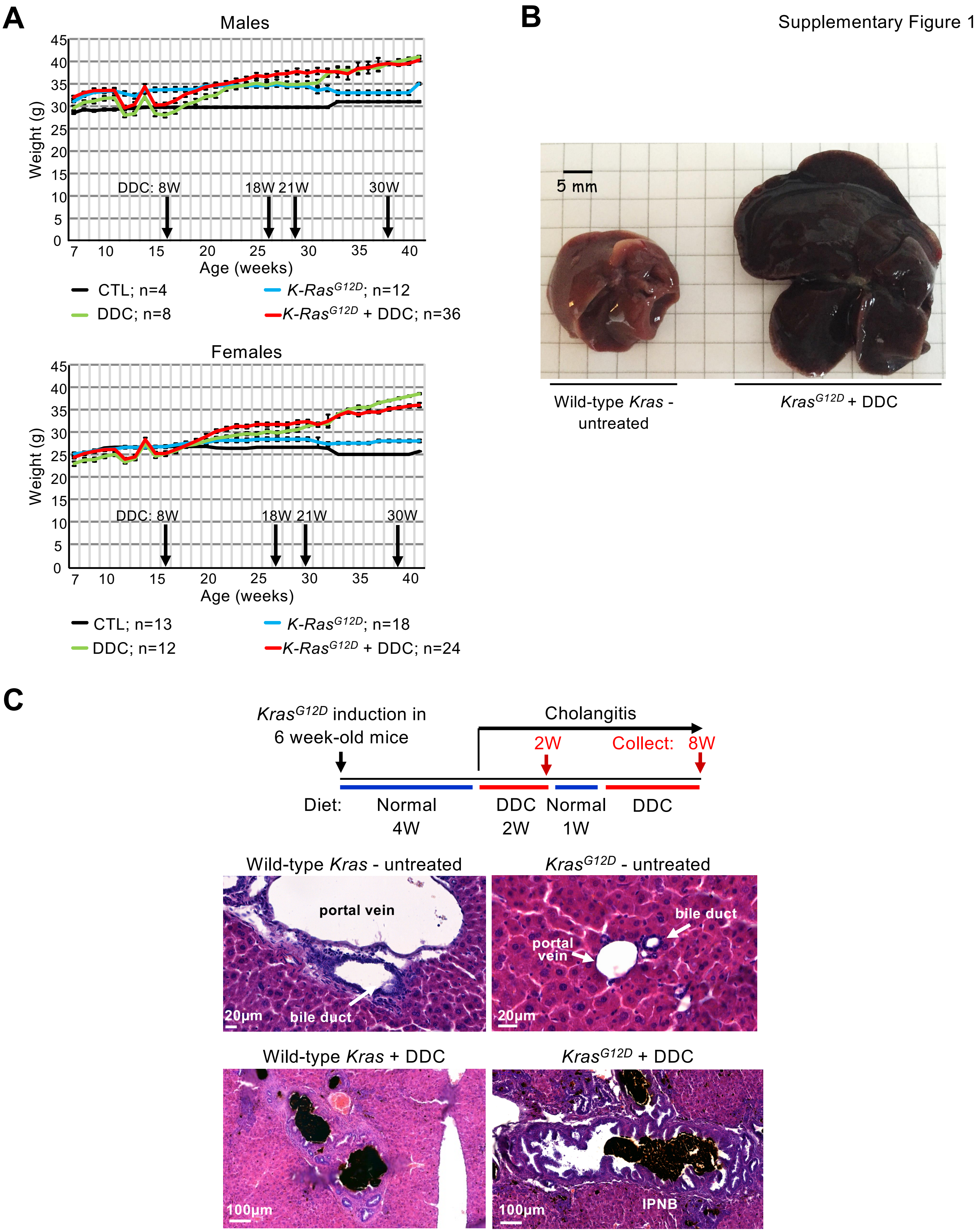

**Supplementary Figure 1**. (*A*) Time-course of mouse body weight progression during DDC treatment. Arrows point to the timing of liver collection after initiation of DDC treatment. Data are means +/- SEM. *(B)* Hepatomegaly in DDC-treated mice. The liver of mice expressing *Kras*$^{G12D}$ and fed a DDC diet for 25 weeks is illustrated. (*C*) Untreated mice expressing *Kras*$^{G12D}$ in cholangiocytes have normal ducts. After 8 weeks of DDC treatment, mice expressing wild type *Kras* showed emerging papillary lesions of large bile ducts, whereas DDC-treated mice expressing *Kras*$^{G12D}$ in cholangiocytes displayed IPNBs.

Supplementary Figure 2

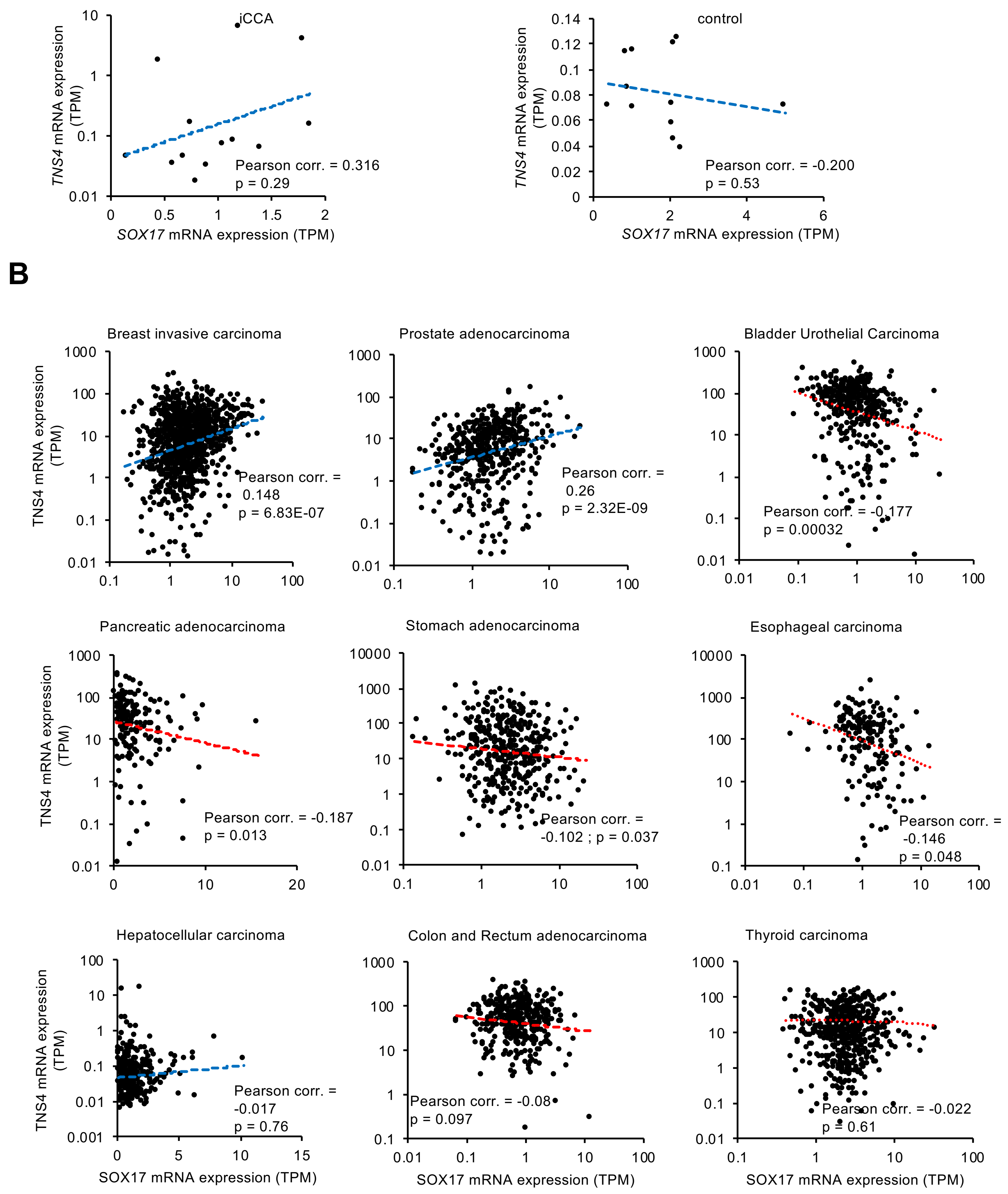

**Supplementary Figure 2.** (*A*) Correlation between *SOX17* and *TNS4* mRNA expression in the GSE119336 cohort. The correlation is not statistically significant in iCCA and the number of samples is low, but clearly differs from the controls. Control is non-tumor adjacent tissue. (*B*) Correlation between *SOX17* and *TNS4* mRNA expression in TCGA cohorts. For each cohort from TCGA, we converted the “scaled_estimate” in the “illuminahiseq_rnaseqv2_unc_edu_Level_3_RSEM_genes” file into TPM by multiplying by $10^6$.

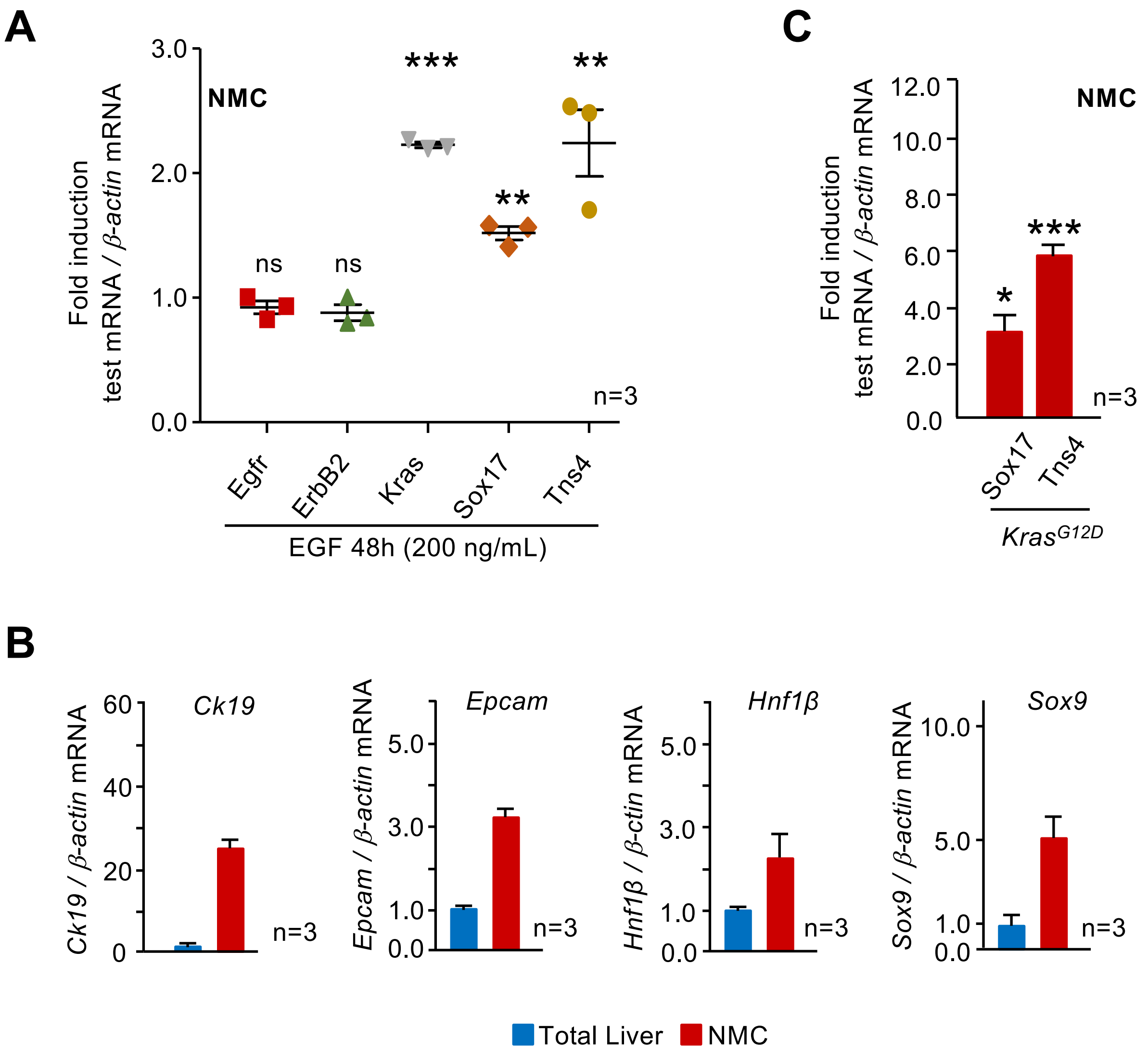

**Supplementary Figure 3.** (*A*) Expression of *Kras*, *Sox17* and *Tns4* is upregulated in NMC cells incubated with EGF, as compared to untreated cells. (*B*) NMC cells express cholangiocyte markers. The level of marker expression in cholangiocytes is compared with that in total adult mouse liver. (*C*) Transient transfection of *Kras*$^{G12D}$ stimulates *Sox17* and *Tns4* expression in NMC cells.

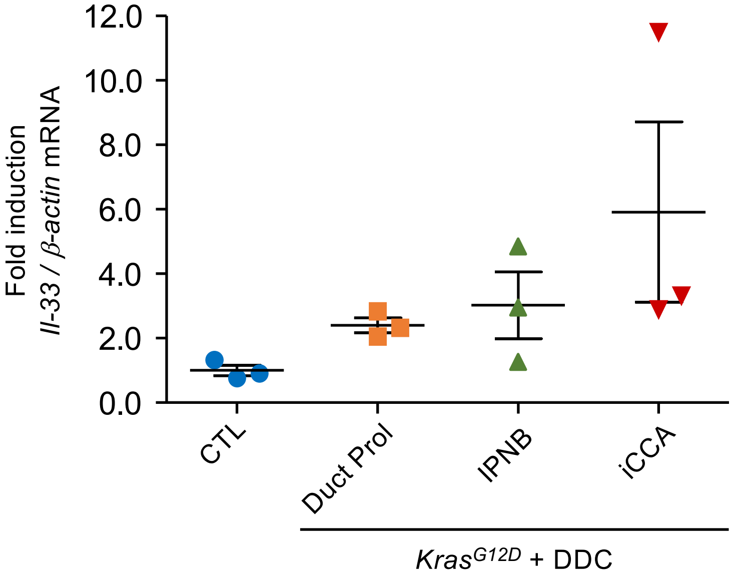

**Supplementary Figure 4.** Expression of IL-33 mRNA in control cholangiocytes purified from tamoxifen-treated Opn-Cre$^{ER}$/Rosa26R$^{eYFP}$ mice and from ductular proliferations (Duct Prol), IPNB and iCCA from Opn-Cre$^{ER}$/LSL-Kras$^{G12D}$ mice treated with tamoxifen to induce Kras$^{G12D}$ and fed a DDC diet for 25 weeks. Data are means of fold inductions +/- SEM. * $p<0.05$, ** $p<0.01$ and *** $p<0.001$.
